# Gene tree conflict between genomic and functional partitions impacts phylogenetic inference in Zygophylloideae

**DOI:** 10.64898/2026.01.16.699981

**Authors:** Pieter de Wet van der Merwe, Michael D. Pirie, Dirk U. Bellstedt

## Abstract

The Zygophyllaceae subfamily Zygophylloideae includes seven genera and around 180 species distributed worldwide particularly in arid areas where they show adaptations to drought including C_4_ and CAM photosynthesis. Phylogenetic relationships of Zygophylloideae have been analysed in several studies using Sanger sequenced markers, but none have been able to resolve all deeper relationships between genera in the subfamily. Here, we use ion semi-conductor sequencing of chloroplast and nuclear ribosomal regions to analyse these relationships and assess why they have proved so challenging to resolve. We sequenced 39 chloroplast genes and nuclear ribosomal 18S, 5.8S, and 26S genes and internal transcribed spacers (ITS) for key species of *Zygophyllum sensu stricto, Roepera, Tetraena, Fagonia*, *Zygophyllum stapffii* and monotypic *Augea capensis*; and *Tribulus* of the subfamily Tribuloideae as outgroup. We aligned these with data from published chloroplast genomes followed by phylogenetic analyses using Maximum Likelihood and Bayesian approaches, summarising phylogenetic results separately for non-coding introns and intergenic spacers, and for genes that are, or are not, involved in photosynthesis. We combined the chloroplast data with sequences of three highly variable noncoding chloroplast regions used in previous studies of greater numbers of taxa, and similarly the nuclear ribosomal data with more densely sampled ITS sequences and assessed whether adding such data resulted in improved support for consistent topologies. We used a chloroplast sequence matrix to infer a dated phylogeny. Adding data did not always improve phylogenetic resolution which was apparently impacted by non-informative variation in chloroplast photosynthesis related genes as well as by conflicting chloroplast and nuclear signals. Chloroplast genes not related to photosynthesis provide robust support for a topology that is consistent with independent data and may therefore represent a useful species tree hypothesis. The cause of the gene tree conflicts remains unclear: potential explanations include functional constraints in photosynthesis related genes and introgressive hybridization and/or incomplete lineage sorting during an ancient rapid divergence. Further testing will require a broader sampling of lineages with different photosynthetic modes and multiple informative, independent, and selectively neutral gene trees.

## Introduction

The flowering plant family Zygophyllaceae has a world-wide distribution (Sheahan and Chase, 1996). They can be trees, such as species of the genus *Balanites* Delile; shrubs such as the well-known creosote bush, *Larrea tridentata* (DC.) Coville, from the USA; or annual herbs, such as *Tribulus terrestris* L.. The family includes examples of tropical rainforest trees in South America, such as *Bulnesia* Gay and *Guaiacum* Plum ex L. which are known for their particularly hard wood. However, Zygophyllaceae are more typically characterised by adaptations to drought tolerance that have allowed the family to radiate into all arid areas of the world.

Sheahan and Chase (1996) presented the earliest molecular systematic studies of the Zygophyllaceae based on sequence data from the chloroplast *rbc*L gene. They reviewed previous morphology-based classifications of the family and re-circumscribed subfamilies Zygophylloideae, Tribuloideae, Seetzenioideae, Larreoideae and Morkillioideae (Sheahan and Chase, 1996). According to their redefinition, the subfamily Zygophylloideae consisted of the genera *Zygophyllum* L. (which included ca. 150 species at that time, referred to here as *Zygophyllum sensu lato*), *Fagonia* L. (30 species) and the monotypic (at that time) genera *Augea* Thunb. and *Tetraena* Maxim. Sheahan and Chase (2000) published a more detailed hypothesis for relationships within the subfamily Zygophylloideae based on chloroplast *trn*LF and *rbc*L sequences of species of *Zygophyllum sensu lato,* of *Fagonia*, and one each of *Augea* and *Tetraena*. Beier et al. (2003) expanded the taxon sampling considerably and, based on non-coding chloroplast *trn*L intron sequences and a morphological character dataset, discovered that *Zygophyllum sensu lato* was polyphyletic. They proposed a new subgeneric classification for Zygophylloideae, recognising a more narrowly circumscribed *Zygophyllum* (referred to hereafter as *Zygophyllum sensu stricto*), resurrecting the genus *Roepera* A.Juss. to include most members of *Zygophyllum* subgenus *Zygophyllotypus* (as recognised by Van Huysteen, 1937), expanding *Tetraena* to include most members of *Zygophyllum* subgenus *Agrophyllum* (as recognised by Van Huysteen, 1937), retaining *Fagonia* and the still monotypic genus *Augea*, and describing a new genus, *Melocarpum* Beier & Thulin. Beier et al. (2004) sequenced the nuclear ribosomal ITS region of all species of *Fagonia* plus Zygophylloideae outgroups to study the relationships and biogeography of *Fagonia*, the only genus in the subfamily to range into the New World. Bellstedt et al. (2008) expanded the taxon sampling of southern African members of the subfamily Zygophylloideae, and, based on a phylogeny derived from *trn*LF and *rbc*L sequences, mostly confirmed the results of Beier et al. (2003). However, they also included *Zygophyllum stapffii* Schinz from the coastal areas of the Namib desert in Namibia and Angola, retrieving this species as sister to *Augea capensis* with high bootstrap support and thereby implying paraphyly of *Zygophyllum s.s.*.

Subsequent studies also based on a limited number of Sanger-sequenced plastid and nuclear ribosomal region sequences have expanded representation of species: Wu et al. (2015) sequenced the *trn*LF and ITS of *Zygophyllum sensu stricto* and a greater sampling of *Tetraena* from Asia and combined these with published sequences of other groups to produce a combined nuclear and chloroplast phylogeny of the subfamily. Wu et al. (2018) sequenced *rbc*L of sixteen additional Asian species of *Zygophyllum sensu stricto* and combined them with *trn*LF and ITS. Lauterbach et al. (2016) focussed on relationships and adaptations to aridity within *Roepera* and *Tetraena* in southern Africa, sequencing the chloroplast *atp*I*-atp*H spacer, *trn*G intron and *trn*LF of an even greater number of taxa. Regardless of molecular and taxon sampling, all of these analyses (Bellstedt et al., 2008, Wu et al., 2015; Lauterbach et al., 2016; Wu et al., 2018) retrieved, with high support, clades corresponding to *Tetraena*; *Roepera*; *Fagonia* plus *Melocarpum* (Engl.) Beier & Thulin (with two *Melocarpum* species sister to *Fagonia*); *Augea capensis* Thunb. plus *Zygophyllum stapffii* / *Z. orbiculatum* Welw.ex Oliv. (the latter treated as conspecific by Bellstedt et al., 2008); and *Zygophyllum sensu stricto*. Relationships between these clades were inconsistent and/or weakly supported in all of these studies.

There are several possible explanations for lack of support for inter-generic relationships in the subfamily Zygophylloideae. First, it could be a lack of sufficient sequence data and/or rapid lineage divergence. Molecular dating analyses have indicated an ancient rapid radiation event between 20 to 13 Mya (Bellstedt et al., 2012) or 38 to 28 Mya (Wu et al., 2015; Wang et al., 2018) which, depending on relaxed clock calibration, occurred c. 38-20 mya, the younger bound corresponding to the onset of global aridification during the start of the Miocene (Bellstedt et al., 2012). Resolving rapid successions of speciation events requires more variable sequence data, and the modest numbers of Sanger sequenced markers used may simply have been insufficient. This deficit could be addressed using high throughput sequencing of larger numbers of chloroplast genes or whole genomes (Barrett et al., 2013, 2014), particularly to establish relationships between groups in ancient rapid radiations (Zhang et al., 2016). Zhang et al. (2021) analysed complete chloroplast genomes of 24 samples of *Zygophyllum sensu stricto* occurring in China plus *Tetraena mongolica* Maxim. as an outgroup. The results were promising, delivering a generally robustly supported topology. However, due to the restricted taxon sampling the unresolved basal nodes in Zygophylloideae were not represented.

Beyond a simple lack of data, other explanations for a failure to infer relationships could involve differences in phylogenetic signal between nuclear, chloroplast and mitochondrial genomes or even within genomes. Such conflict could be the result of reticulate processes such as hybridization or demographic processes resulting in incomplete lineage sorting (De Queiroz et al., 1995; Maddison, 1997). In both cases differences between gene trees can be meaningful for interpretation of evolutionary history, but they make inferring a species tree more challenging (Nakhleh, 2013) and require more sequence data for effective comparison of individually resolved, independent gene trees, e.g. from targeted high throughput sequencing (Johnson et al, 2019; Larridon et al., 2020; Morales Briones et al., 2021; Yardeni et al., 2022).

Whether derived from Sanger or high throughput sequencing, functional constraints or even convergence could limit or even undermine the usefulness of particular genes (Shen et al., 2017). In the case of Zygophylloideae, those encoding proteins involved in the different modes of photosynthesis could be such an example. C_4_ photosynthesis has been inferred to have evolved from C_3_ ancestors numerous times independently in flowering plants in response to the reduction in atmospheric carbon dioxide (CO_2_) from the Oligocene onwards (Pagani et al., 2005; Christin et al., 2011). This photosynthesis type has the advantage that atmospheric CO_2_ can be captured in chemical intermediates at night, enabling the plant to close its stomata during the day avoiding moisture loss. C_4_ plants can therefore survive better in arid environments. C_4_ photosynthesis evolved in the subfamily Zygophylloideae in *Tetraena simplex* (L.) Beier & Thulin and independently in clades of *Tribulus* L., *Tribulopsis* F.Muell.and *Kallstroemia* Scop. species in the sister subfamily Tribuloideae (Lauterbach et al., 2019). Lauterbach et al. (2016) analysed adaptations in leaf morphology in the genera *Tetraena* and *Roepera* showing that the vascular bundles which typically facilitate C_4_ photosynthesis only evolved in *Tetraena simplex* and not in other species in the genus or in *Roepera*. However, diverse other leaf anatomical types in *Tetraena* do suggest similar adaptations in other species. The genes encoding for the proteins that lead to C_4_ photosynthesis involves many genes encoded in the chloroplast genome such as the *rbc*L gene (Christin et al., 2008), but also in genes that control leaf structure which therefore must be encoded in the nuclear genome. The *rbc*L gene has been used extensively in phylogenetic reconstruction at all hierarchical levels in plant phylogenetics (Chase et al., 1993; APG I., 1998; APG II, 2003, APG III, 2009; APG IV, 2016). However, it is known to exhibit specific convergent mutations in C_4_ lineages (Christin et al., 2008).

Another photosynthetic mechanism that serves as an adaptation for aridity, Crassulacean acid metabolism (CAM), evolved in *Roepera cordifolia* L.f. Beier & Thulin (Matimati et al. 2012). It is not known whether CAM is present in other species of *Roepera*. However, CAM photosynthesis is also present in the New-World species *Bulnesia retama* (Gillies ex Hook. & Arn.) Griseb. in the subfamily Larreoideae (Mok et al. 2023). CAM metabolism involves a third, very different photosynthetic mechanism for photosynthetic processes to occur in plants exposed to aridity and is not facilitated by a Kranz type morphology as in C_4_ plants but rather an intracellular adaptation. Evolution towards this type of photosynthesis apparently requires mutation and expression of a different repertoire of genes, including *rbc*L, to those in plants that possess C_3_ or C_4_ photosynthetic mechanisms (Berry et al., 2013). It is important to test whether the rbcL gene and potentially other genes in the chloroplast genome might show convergent signals that impact phylogenetic inference in plant groups in which either C_3_, C_4_ or CAM photosynthesis occurs.

Here, we analyse numerous coding and non-coding genes from both the chloroplast and nuclear genomes to try to resolve intergeneric relationships in the subfamily Zygophylloideae which have not been resolved in any studies in the subfamily before. We focus on a limited number of key species in the major lineages in subfamily of *Zygophyllum sensu stricto, Roepera, Tetraena, Fagonia*, and the possibly monotypic lineage of *Zygophyllum stapffii/orbiculatum* and the monotypic *Augea*, with a representative member of the subfamily *Tribuloideae* as outgroup. We compared phylogenies generated from sequencing a large part of the chloroplast genome, subsets thereof (photosynthetic genes versus non-photosynthetic genes) and sequences of a smaller number of highly variable chloroplast genes with much denser sampling of taxa. Furthermore, we compared phylogenies generated from the entire nuclear ITS region of key taxa to those generated by the addition of ITS sequences with much denser taxon sampling. Finally, we compared chloroplast and nuclear phylogenies.

## Materials and Methods

### Chloroplast isolation, DNA purification and ion semi-conductor sequencing

Fresh leaf material was obtained from selected species in six genera in the subfamily Zygophylloideae: *Roepera foetida*, *Zygophyllum fabago*, *Zygophyllum stapffii, Tetraena turbinata* ined., *Augea capensis*, *Fagonia rangei* and from an outgroup *Tribulus terrestris* (see Supplementary Table 1). Intact chloroplasts were isolated using a Chloroplast Isolation Kit from Sigma-Aldrich. Approximately 30 g of fresh plant leaf material was macerated in a blender using the detailed instructions of the kit. A 40%/80% Percoll gradient was used to collect intact chloroplasts at the interphase of the two gradients using a Pasteur pipette. After subsequent steps the final isolated chloroplast pellet was resuspended in approximately 500 µl Chloroplast Isolation Buffer (CIB) supplied with the isolation kit. This suspension was used in an adapted protocol of Doyle & Doyle, 1987 where upon 500 µl of 2XCTAB was added to lyse the intact chloroplasts without a grinding step, releasing the DNA into solution. The protocol was then followed as described. Contaminating RNA was removed using RNA digestion. The concentration of the purified DNA was determined using a NanoDrop 1000 Spectrophotometer and ion semi-conductor sequencing was performed at Central Analytical Facility (CAF), University of Stellenbosch following the manufacturer’s instructions. The sequence reads generated by the ion semi-conductor sequencing were quality filtered according to the following criteria: primer dimers, identical sequences reads, and reads that were less than 25 bp long were removed. Furthermore, quality trimming of reads was performed from the 3’ end using a sliding window of 30 and a Q20 threshold.

### Chloroplast gene contiguous sequence assembly and alignments

The quality filtered short sequence reads obtained from the ion semi-conductor sequencing were subjected to *de novo* assembly to generate contiguous sequences (contigs) using either Newbler (http://454.com/products/analysis-software/index.asp) or Mira (http://sourceforge.net/projects/mira-assembler/) using default settings and were converted to fasta format. Contiguous sequences were generated from the chloroplast, the nuclear and the mitochondrial genomes.

Annotation of the generated contigs was performed in Geneious Prime 2023.1.2 using the complete *Corynocarpus laevigatus* J.R.Forst. & G. Forst. chloroplast genome as a reference genome (NC_014807) (Atherton et al., 2010). This allowed for the rapid identification of chloroplast contigs. Only the 41 chloroplast genes that were identified in all seven sequenced taxa were retained to exclude missing data. Nine additional whole chloroplast genomes in the Zygophyllales were downloaded from GenBank. These include two specimens from the Krameriaceae; *Krameria lanceolata* O. Berg (NC_043801) and *Krameria bicolor* S. Watson (NC_043800), (Gonçalves et al., 2019), and seven species from the family Zygophyllaceae; *Tribulus terrestris* (MK341055) (Xi et al., 2014) in the family subfamily Tribuloideae; *Bulnesia arborea* (Jacq.) Engl. (see Supplementary Table 2) (Moore et al., 2010), *Guaiacum angustifolium* Engelm. (NC_043796) (Gonçalves et al., 2019) and *Larrea tridentata* (KT272174) (Countway et al., 2015) in the subfamily Larreoideae; and *Fagonia cretica* L. (OM952137) (Sonchhatra et al., 2022), *Tetraena mongolica* (ON408223) (Yang, 2022) and *Zygophyllum xanthoxylum* (Bunge) Maxim. (NC_052769) (Xu et al. 2021). From these nine complete chloroplast genomes the 41 abovementioned genes were identified and aligned with the sequences retained from the ion semi-conductor sequencing described above. The species of which the sequences were downloaded from Genbank were selected so that the genera *Fagonia*, *Tetraena* and *Zygophyllum* s.s. could each be represented by their two most distantly related species. These species were targeted to ensure that the phylogenetic analyses performed on the combined sequences would represent the oldest nodes in each of the respective genera i.e. *Fagonia cretica* and *Fagonia rangei* Loes. Ex Engl. (Beier et al., 2004); *Tetraena turbinatum* ined. and *Tetraena mongolica* (Lauterbach et al., 2016) and *Zygophyllum fabago* and *Zygophyllum xanthoxylum* (Lauterbach et al., 2016) respectively.

Further inspection of the 41 gene sequence alignments revealed that two were non-coding gene regions (*trn*A and *trn*I contain large introns) and that three of the remaining 39 coding genes contained introns (*pet*B, *pet*D, *rpo*C1). These introns were spliced out of the coding genes to not disrupt the reading frame of the protein-coding genes, but the three introns were retained and added to the other two noncoding regions (the abovementioned large *trn*A and *trn*I introns) to give an alignment matrix of 5 noncoding regions (referred to as 5NC). The alignment matrix of remaining 39 coding genes (referred to as 39C) was then further subdivided into those genes directly involved in the photosynthetic process, totaling 21 (referred to as 21PG), and those genes that were not directly involved in the photosynthetic process (referred to as 18NPG) (see Supplementary Table 2). All of these alignment matrices contained the sequences of 16 taxa (9 ingroup [subfamily Zygophylloideae] and 7 outgroup [2 most distant in the family Krameriaceae, 2 in the subfamily Tribuloideae, and 3 in subfamily Larreoideae] and were complete, i.e. they contained no missing data.

To augment the non-coding gene sequence information in the phylogenetic analyses, three highly variable noncoding sequences (referred to as 3HVNC) in the chloroplast genome, i.e*. trn*LF *(trn*L *intron + trn*LF *spacer)*, *trn*G and *atp*I*-atp*H, were obtained from various sources. Firstly, one each per monophyletic species but including the northern and southern populations of *Tetraena simplex* and *Tetraena decumbens* (Delile) Beier & Thulin respectively, were obtained from previous studies (Sheahan and Chase, 1996 and 2000, Bellstedt et al., 2008, Lauterbach et al., 2016). Secondly, those obtained from the contiguous sequences generated by ion semi-conductor sequencing generated in this study and thirdly, those obtained from the whole genome sequences downloaded from GenBank. They were combined in Geneious Prime 2023.1.2, aligned using MUSCLE v3.8.31 (Katoh & Standley, 2013) followed by manual refinement to generate a large chloroplast gene alignment matrix consisting of 121 taxa. The sequences of the 3HVNC regions could not be utilised from the whole genome sequence of *Fagonia cretica* (OM952137) because of the poor quality of the genome assembly deposited on GenBank. In previous studies it was found that the *trn*L*-trn*F spacer region of the *trn*LF sequence region of *Augea capensis* could not be sequenced with standard primers (Taberlet et al., 1991). Ion semi-conductor sequencing revealed that the *trn*F gene was inverted in *Augea capensis*. This gene sequence was then reverse-complemented and incorporated into the *trn*L*-trn*F alignment matrix. To study the effect of the addition of the 3HVNC sequences on the phylogenetic analyses, an alignment matrix consisting of these 3HVNC sequences of 15 of the 16 taxa (as those of *Fagonia cretica* were missing) of which the 39 coding genes and 6 noncoding genes were sequenced or obtained from GenBank, was generated. Alignments of increasing numbers of the 3HVNC sequences of an evenly distributed number of species in the different genera were then generated (34, 96 and 121 taxa) to study the effect of taxon addition to the 16-taxon coding and noncoding gene matrices on the phylogenies resulting from their combined phylogenetic analyses.

### Taxon sampling, DNA extraction and nuclear ITS cassette gene sequencing and alignment

In order to obtain nuclear gene sequences from the taxa from which chloroplast sequences were generated to make comparisons to nuclear phylogenies, an attempt was made to retrieve nuclear sequences from the quality filtered ion semi-conductor sequencing data. It was found that the complete nuclear ITS cassette region consisting of the 18S rRNA, ITS1, 5.8S rRNA, ITS2 and 26S rDNA gene cassette region of the seven taxa from which chloroplast sequences were generated could be retrieved from the Ion semi-conductor sequencing data due to nuclear DNA contamination of the chloroplast enriched fraction. However, in order to generate comparable phylogenies, the nuclear ITS cassette sequences of appropriate outgroups were required. The complete ITS cassette sequences were not available for *Bulnesia arborea* and several gaps were identified in the ITS cassette region sequences of *Tetraena turbinata* ined. and of *Tribulus terrestris*. For this reason, silica gel dried leaf material of *Bulnesia arborea* was obtained from the Royal Botanical Garden Edinburgh, UK and DNA was extracted from it following Doyle and Doyle (1987).

The complete ITS cassette region was amplified using specifically designed primers (primers are listed in Supplementary Table 3) from the DNA isolated from *Bulnesia arborea*. PCR products were purified and sequenced using standard Sanger sequencing (CAF, Stellenbosch University). Some of the primers generated for the amplification of the whole ITS region of the *Bulnesia arborea* could be used to fill in the missing regions in the assembled contigs of *Tribulus terrestris* and *Tetraena turbinata* ined. via Sanger sequencing. The generated contiguous sequences were then used to construct a complete nuclear 18S rDNA-ITS1-5.8S rDNA-ITS2-26S rDNA cassette sequence matrix of a total of 8 taxa.

### ITS gene sequencing and alignment

Samples of some additional taxa in the Zygophylloideae were obtained from the Royal Botanic Gardens, Kew, UK, (see Supplementary Table 4) and DNA was isolated as described above. Amplification of the ITS region (ITS1, 5.8s RNA and the ITS2) of these taxa was achieved using primers 17SE and 26SE of Sun et al. (1994), also referred to as AB101 and AB102 by Douzery et al. (1999) (primers are listed in Supplementary Table 3). Alternatively, the primers P16 (forward, TCA CTG AAC CTT ATC ATT TAG AGG A), P17 (forward, CTA CCG ATT GAA TGG TCC GGT GAA), P25 (reverse, GGG TAG TCC CGC CTG ACC TG) 26S-82R (reverse, TCC CGG TTC GCT CGC CGT TAC TA) and 2G (reverse, GTG ACG CCC AGG CAG ACG T) were used to successfully amplify the gene region. PCR reactions were performed in an Applied Biosystems Veriti 96 well Thermal Cycler in 25 µl reactions. Each tube contained 2.5 mM MgCl_2_, 1x JMR-455 buffer (Southern Cross Biotechnology, Cape Town, RSA), 1 U of Super-Therm Taq polymerase (Southern Cross Biotechnology, Cape Town, RSA), 200 µM of each dNTP and 0.5 µM of the forward and reverse primer respectively. Amplification was achieved using 35 cycles of 1 min at 94 °C, 1 min at 55 °C, 90 s at 72 °C, and final extension of 6 min at 72 °C.

PCR products were sequenced with the primers as specified above at Stellenbosch University’s Central Analytical Facility. Forward and Reverse sequencing chromatograms were superimposed and base calling corrected in ChromasPro v2.1.8 to generated consensus sequences.

Consensus sequences were imported and aligned in Geneious Prime 2023.1.2 using MUSCLE v3.8.31 (Katoh & Standley, 2013). Manual adjustments were made to ensure correct alignment. Indels were removed and treated as missing data. The sequence matrix therefore consisted of the ITS1, 5.8s RNA and the ITS2 region of 59 taxa. The ITS1, 5.8s RNA and the ITS2 region of the 8 taxa from which the full ITS cassette region was sequenced was then added to the alignment matrix of 59 taxa to give a total ITS alignment matrix of 67 taxa.

### Strategies for phylogenetic analyses of sequence alignments

The following aligned matrices were analysed using the methods of phylogenetic analysis described hereafter:

1. a. The coding gene sequences (39C, 16 taxa) and subdivisions thereof into a non-photosynthesis coding gene matrix (18NPG, 16 taxa) and a photosynthesis coding gene matrix (21PG, 16 taxa). b. A 5NC non-coding gene matrix of 16 taxa combined with a 3HVNC matrix in combinations of increasing taxon numbers (16, 34, 96 and 122). c. In order to establish the effect of adding non-coding sequence data (1b, 16 taxa) to the coding sequence alignment matrices 39C and 18NPG (1a, 16 taxa), combinations of these alignment matrices were made: Coding non-photosynthetic and non-coding sequences (18NPG + 5NC +3HVNC, 16 taxa), all coding and non-coding sequences (39C + 5NC +3HVNC, 16 taxa) d. In order to establish the effect of adding non-coding sequence data of additional taxa (1b, 34, 96 and 121 taxa) to the coding sequence alignment matrices 39C and 18NPG (1a, 16 taxa) combinations of these alignment matrices were made: Coding and non-coding sequences of 16 taxa with increasing numbers of the 3HVNC sequences: ((39C + 5NC, 16 taxa) +(3HVNC of 34 taxa)); ((39C + 5NC, 16 taxa) +(3HVNC of 96 taxa)); ((39C + 5NC) + (3HVNC of 121 taxa). 3HVNC were not available for *Fagonia cretica* from Genbank although its coding gene sequences were available.
2. a. The complete ITS cassette dataset (18S rRNA, ITS1, 5.8S rRNA, ITS2 and 26S rRNA) consisting of the sequences of the 6 Zygophylloideae ingroup taxa and *Bulnesia arborea* and *Tribulus terrestris* as outgroups. b. The ITS data only (ITS1, 5.8SrRNA and ITS2) of 67 taxa listed in Supplementary Table 3. c. The combination of 2a and 2b representing the same 67 taxa.

A summarizing diagram of the aligned matrices and their phylogenetic analyses, together with the figure numbers in which their results will be presented is shown in Figure 1.

**Figure 1:**
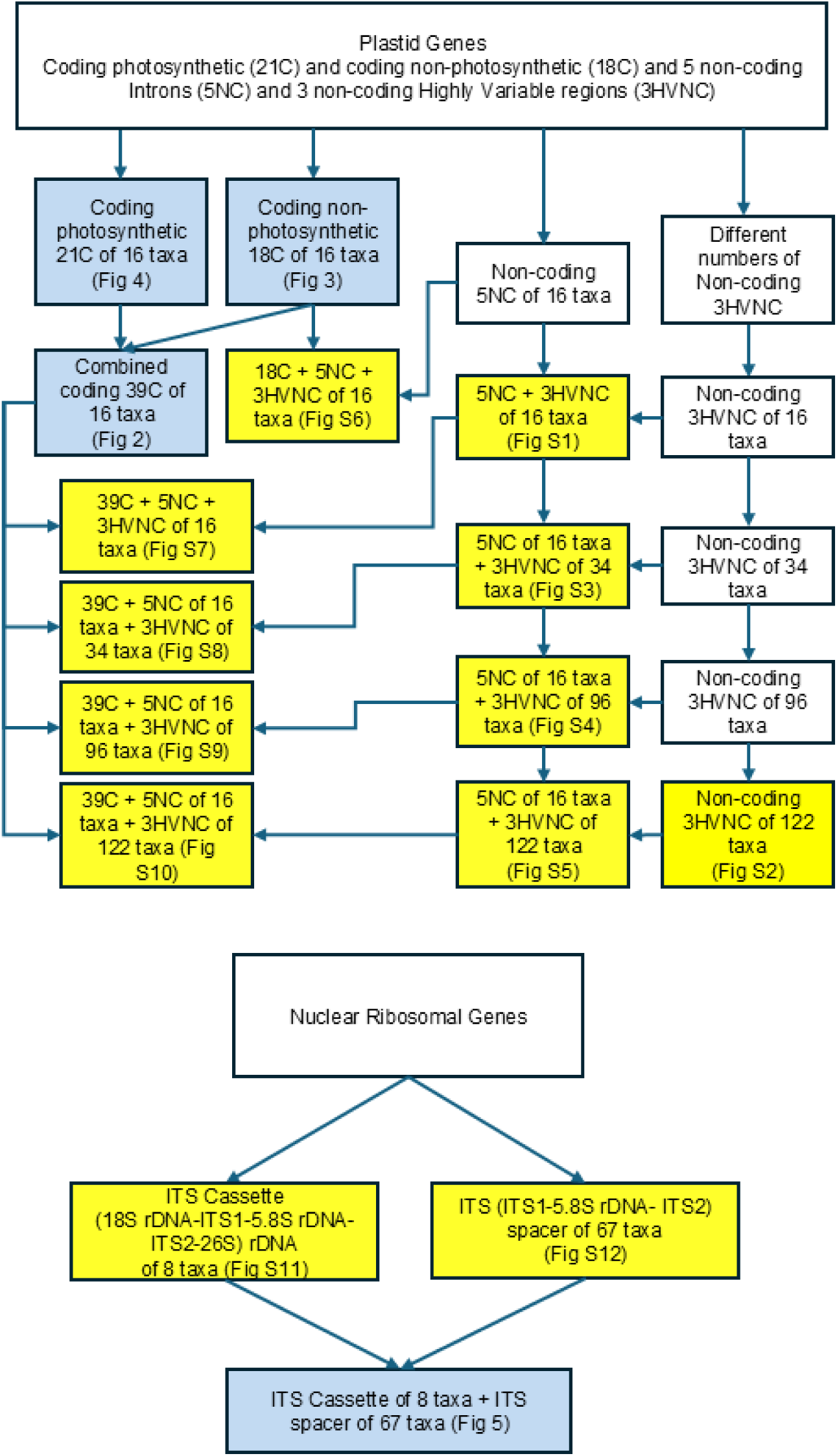
A diagram summarising the aligned matrices generated in this study and their phylogenetic analyses. The matrices of which the phylogenetic analyses are presented as figures and supplementary figures are highlighted in light blue and yellow respectively. The figures and supplementary figure numbers as presented in the results are shown in brackets.

### Phylogenetic analyses

All individual sequence alignments were analysed to infer the best-fitting evolutionary model using jmodeltest v2.1.6 as implemented on the CIPRES Science Gateway (Miller et al. 2010). In all cases the best-fitting model was found to be GTR+G. We did not partition the alignment matrices for the phylogenetic analyses.

Alignments of the concatenated chloroplast genes and of the concatenated nuclear ITS region genes as described above were used for phylogenetic analyses. RAxML 8.0 was used for maximum likelihood analyses (Stamatakis, 2014). The most likely trees and bootstrap values were generated. The autoFC function was used to stop generating bootstrap replicates once the support values had stabilised. Alignments comprising only 16 taxa were also analysed using MrBayes 3.2.6 for phylogenetic inference (BI, MrBayes 3.2.6, Ronquist and Huelsenbeck, 2003) on the CIPRES Science Gateway (Miller et al. 2010). Not all alignments were analysed under Bayesian inference because it would have been considerably more time consuming due to the size of the datasets. In all Bayesian analyses, three heated chains and one cold chain were employed, with runs initiated from random trees. Two independent runs were conducted with 10 000 000 generations per run and trees and parameters were sampled every 1 000 generations. Convergence was checked using Tracer 1.5 (Rambaut and Drummond, 2007). For each run, the first 10% of sampled trees were discarded as burn-in. Majority rule consensus trees were generated from the individual trees retrieved from the Bayesian analyses. Tree files were imported into Figtree (Rambaut, 2009) for tree drawing (http://tree.bio.ed.ac.uk/software/figtree/).

### Tests for selection of chloroplast coding gene matrices

All codon aligned protein-coding genes were imported as FASTA files into MEGA v12.1.2 (Kumar et al., 2024) and tested for selection using the “Codon-based Z-test of Selection” function. All three test hypotheses (Neutrality, Positive and Purifying) were tested for using the “Kumar method (Kimura 2-para)” Model/Method. All other settings were left at default.

### Molecular dating analysis

A 15 (rather than 16) taxon 18NPG alignment matrix was used to date the Zygophylloideae phylogeny using BEAST (Drummond & Rambaut, 2007) v1.8.4. One *Krameria* outgroup was removed leaving a single outgroup as required using BEAST. The average age of the node between the Krameriaceae and the Zygophyllaceae of 72.2 Mya (Harris & Davies, 2016) as indicated on TimeTree (Kumar et al., 2022) was used to calibrate the phylogeny, which was dated using an uncorrelated relaxed clock, a lognormal distribution setting and the birth-death model with a standard deviation of 10%. The GTR+G model was also implemented. Two independent analyses were performed for 10 000 000 generations each, sampling every 1 000 generations. Convergence and mixing of the Markov chains was assessed for two independent analyses in Tracer v1.5, where all parameter traces were inspected visually and effective sample sizes (ESS) were confirmed to exceed recommended thresholds after discarding the initial 10% of samples as burn-in. Subsequently a maximum clade credibility tree was generated using TreeAnnotator v1.8.4 from the two combined tree datasets. A maximum clade credibility tree was subsequently generated in TreeAnnotator v1.8.4 from the two combined tree datasets. The dated phylogenetic tree was visualised in FigTree v1.4.2.

## Results

### Chloroplast isolation and chloroplast DNA enrichment

The chloroplast isolation protocol was successful in isolating and enriching DNA from the selected taxa, as was apparent from the subsequently generated ion semiconductor sequence data. However, although the Percoll gradient chloroplast isolation procedure was aimed at the separation of intact chloroplasts from the rest of the cellular content of the plant cells with a view to generating chloroplast DNA sequences only, the generation of some mitochondrial and nuclear sequences from the isolated DNA did indicate contamination with mitochondria and nuclei. However, this apparently had no negative effect on the sequencing of chloroplast DNA and additionally allowed the sequencing of the complete ITS cassette region.

### NGS data

All raw sequence read data of the seven species analysed in this project were submitted to NCBI and are available under PRJNA1526063 (https://www.ncbi.nlm.nih.gov/bioproject/?term=prjna1526063). All *de novo* assembled contiguous chloroplast, nuclear and mitochondrial sequences were deposited in the Stellenbosch University data repository (https://doi.org/10.25413/sun.31238719). Coverage of chloroplast intergenic regions and of mitochondrial sequences in general was too poor to allow complete chloroplast or mitochondrial genome construction with a high degree of accuracy. However, the data did allow the consistent retrieval of 39 chloroplast coding genes (39C) and five chloroplast noncoding gene regions (5NC) from all seven taxa (six ingroup, one outgroup) which were combined with sequences of the same genes obtained from the whole chloroplast genome of three additional ingroup and six outgroup taxa from GenBank to give gene sequences of nine ingroup and seven outgroup taxa.

Alignments of these genes were then used for subsequent phylogenetic analyses. The sequences of the 39 chloroplast coding genes of the seven taxa were deposited on Genbank. Accession numbers of these sequences are shown in Supplementary Table 2.

The characteristics of all alignments including lengths and variability are shown in Table 1 with the topologies of the phylogenies resulting from their phylogenetic analyses.

**Table 1:** A summary of sequence variation of the different gene regions that were sequenced and the topology of the phylogenies generated from them.

| **Informative character and phylogenetic analysis summary** |  |  |  |  |  |  |  |  |
| --- | --- | --- | --- | --- | --- | --- | --- | --- |
| Alignment matrix | Total characters | Constant | Uninformative | Informative | Variable | % variability | Figure | Topology |
| **Chloroplast coding** |  |  |  |  |  |  |  |  |
| 16 taxa Combined 39C | 37764 | 30789 | 1729 | 5246 | 6975 | 18.47 | 2 | F(Z(T(RSA))) |
| 16 taxa Non-Photosynthesis 18C genes | 20253 | 16194 | 1033 | 3026 | 4059 | 20.04 | 3 | F(Z(T(RSA))) |
| 16 taxa Photosynthesis 21C genes | 17511 | 14595 | 696 | 2220 | 2916 | 16.65 | 4 | T(RF)((SA)Z) |
| **Chloroplast non-coding** |  |  |  |  |  |  |  |  |
| 16 taxa non-coding 6NC & 16 taxa 3HVNC | 9422 | 6875 | 743 | 1804 | 2547 | 27.03 | S1 | T(Z(F(RSA))) |
| 121 taxa, non-coding 3HVNC | 4162 | 2689 | 423 | 1050 | 1473 | 35.39 | S2 | T(Z(F(RSA))) |
| 16 taxa non-coding 6NC & 34 taxa 3HVNC | 9422 | 6794 | 802 | 1826 | 2628 | 27.89 | S3 | T(Z(F(RSA))) |
| 16 taxa non-coding 6NC & 96 taxa 3HVNC | 9422 | 6645 | 843 | 1934 | 2777 | 29.47 | S4 | T(Z(F(RSA))) |
| 16 taxa non-coding 6NC & 121 taxa 3HVNC | 9422 | 6606 | 858 | 1958 | 2816 | 29.89 | S5 | T(Z(F(RSA))) |
| **Chloroplast coding & non-coding combinations** |  |  |  |  |  |  |  |  |
| 16 taxa coding Non-Photo 18C & 16 taxa non-coding 6 NC & 3HVNC | 29675 | 23133 | 1829 | 4713 | 6542 | 22.05 | S6 | F(Z(T(RSA))) |
| 16 taxa coding Combined 39C & 16 taxa non-coding 6NC & 3HVNC | 47186 | 37728 | 2525 | 6933 | 9458 | 20.04 | S7 | F(Z(T(RSA))) |
| 16 taxa coding Combined 39C & 16 taxa non-coding 6NC & 34 taxa non-coding 3HVNC | 47186 | 37583 | 2531 | 7072 | 9603 | 20.35 | S8 | T((FZ)(RSA)) |
| 16 taxa coding combined 39C & 16 taxa non-coding 6NC & 96 taxa non-coding 3HVNC | 47186 | 37434 | 2572 | 7180 | 9752 | 20.67 | S9 | F((ZT)(RSA)) |
| 16 taxa coding combined 39C & 16 taxa non-coding 6NC & 122 taxa non-coding 3HVNC | 47186 | 37395 | 2587 | 7204 | 9791 | 20.75 | S10 | F((ZT)(RSA)) |
| **Nuclear coding** |  |  |  |  |  |  |  |  |
| 8 taxa ITS cassette (18S & ITS1 & 5.8S & ITS2 & 26S) | 5930 | 5204 | 422 | 284 | 706 | 11.9 | S11 | (Z(FT))(RSA) |
| **Nuclear non-coding** |  |  |  |  |  |  |  |  |
| 67 taxa ITS (ITS1 & 5.8S & ITS2) | 819 | 365 | 96 | 358 | 454 | 55.4 | S12 | (T(FZ))(RSA) |
| **Nuclear coding & non-coding combined** |  |  |  |  |  |  |  |  |
| 8 taxa ITS cassette & 59 taxa ITS | 5930 | 5073 | 351 | 506 | 857 | 14.4 | 5 | (T(FZ))(RSA) |

### Phylogenetic analyses of combined chloroplast gene sequences

The most likely phylogenetic tree from the maximum likelihood analysis of the 16 taxa 39 coding genes is shown in Figure 2 A. The topology is the same as that of the majority-rule consensus tree from Bayesian inference. posterior probabilities from which are therefore also indicated on the nodes in Figure 2 A. The monophyly of the subfamily Zygophylloideae and the relationships within the Zygophylloideae are strongly supported except for two nodes which were poorly supported in the maximum likelihood analysis (bootstrap support [BS] of 6 and of 41) but more strongly in the Bayesian analysis (posterior probabilities [PP] of 100 and 68 for the corresponding nodes). Where two species represented a genus, they always grouped together with high support. The relationships between *Augea*, *Zygophyllum orbiculatum*/*stapffii* and *Roepera* were strongly supported. Although these represent morphologically distinct genera they have all evolved in Southern African from a common ancestor (Bellstedt et al., 2012) although *Roepera* has subsequently dispersed to Australia and evolved into about 30 species there. Internal branches between the genera were short, whilst the terminal branches were much longer.

**Figure 2.**
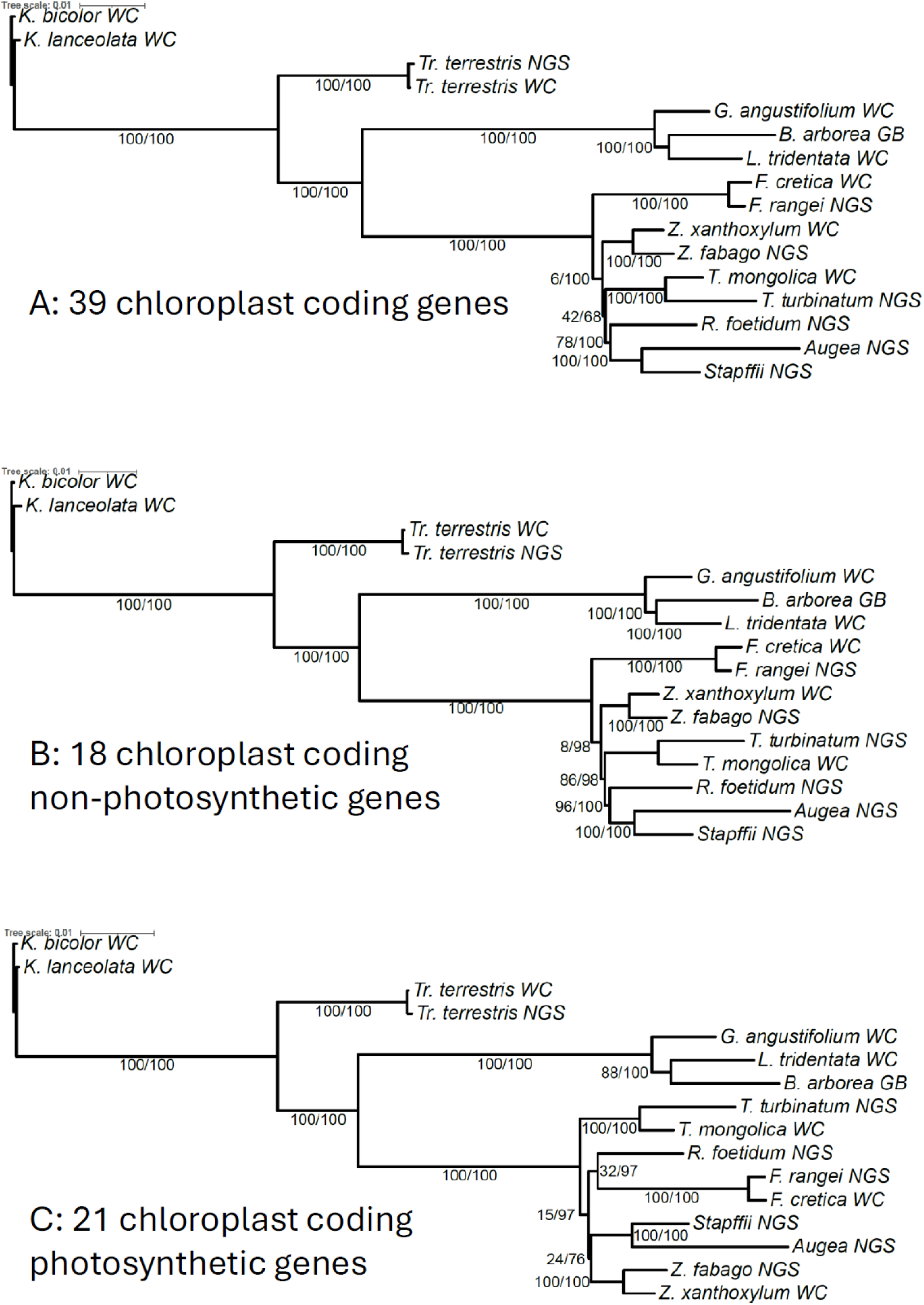
A: The most likely phylogenetic tree from the maximum likelihood analysis of the 16 taxa 39 chloroplast coding genes alignment matrix. B: The most likely phylogenetic tree from the maximum likelihood analysis of the 16 taxa 18 non-photosynthetic coding genes alignment matrix. C: The most likely phylogenetic tree from the maximum likelihood analysis of the 16 taxa 21 photosynthetic coding genes alignment matrix. Maximum likelihood bootstrap values and Bayesian posterior probabilities are indicated on the nodes. The genus names of taxa are abbreviated to a single letter; *Z*. for *Zygophyllum*; *T*. for *Tetraena*, and *R*. for *Roepera*, but the monotypic genus *Augea* was not abbreviated. *Zygophyllum stapffii*/*orbiculatum* is indicated as ‘Stapffii’. Those taxa of which sequences were generated in this study, were indicated by the suffix NGS (next generation sequencing), and those taxa of which the sequences had been obtained from whole chloroplast sequences deposited on GenBank are indicated with suffix WC or in the case of *Bulnesia arborea,* GB.

To test potential conflict within the dataset the 16 taxa 39C gene matrix was subdivided into the 18NPG gene matrix consisting of genes not directly involved in the photosynthetic process and the 21PG gene matrix consisting of genes involved in the photosynthetic process. Phylogenies inferred from these matrices are shown in Figures 2 B and 2 C respectively. The phylogeny generated using 18NPG gene matrix shows the same topology as the phylogeny generated from the 39C gene matrix but gives a better-supported topology compared to the 39C matrix (maximum likelihood BS ≥70 and Bayesian PP ≥95, except for the basal node between genus *Fagonia* and the rest of the subfamily in the maximum likelihood phylogeny (ML, 8%BS, Bayes, 98%BS), as shown in Figure 2 B. By contrast, the 21PG gene matrix consisting of genes involved in the photosynthetic process shows poor support of three nodes (See Figure 2 C). Additionally, the topology of the 21PG phylogeny differed from the 39C and the 18NPG phylogenies and has three nodes that are poorly supported (3 ML nodes < 50%BS, 1 Bayes node <95%BS support).

The phylogenetic tree inferred from the 16 taxa eight non-coding sequences (5NC and 3HVNC) is shown in Figure S1 and that of the 3HVNC sequences only in supplementary Figure S2. Both failed to resolve the basal intergeneric relationships (see Table 1). The addition of increasing numbers of 3HVNC sequences (representing 34, 96 and 121 taxa, Supplementary Figures S3, S4 and S5) to the 16 taxa 5NC gene sequence matrix, resulted in weakened support values. None resulted in a phylogeny of which the basal intergeneric relationships were fully resolved and supported.

Phylogenetic results of combined 16 taxa 18NPG gene matrix and 16 taxa 5NC and 3 HVNC gene matrices are presented in Figure S6 and that of the 16 taxa 39C coding gene matrix with the 16 taxa 5NC and 3 HVNC alignments in Figure S7. Both the addition of the 16 taxa noncoding genes (5NC and 3HVNC) as well as the 16 taxa 21PG gene matrix consisting of genes directly involved in the photosynthetic process, weakened the support compared to that of the 16 taxa 18NPG gene matrix consisting of genes not directly involved in the photosynthetic process. The 18NPG 16 taxa plus noncoding genes (5NC and 3HVNC, supplementary Figure S6) showed even weaker support of the basal node (ML, 2%BS; Bayes, 83%BS) and in the 39C 16 taxa plus noncoding genes (5NC and 3HVNC, supplementary Figure S7) the basal node support dropped (to ML, 0%BS; Bayes, 99%BS) and the second most basal node dropped as well (to ML, 72%BS; Bayes, 98%BS).

Additionally, as more sequences of the 3HVNC were added to the 16 taxa 39C and 5NC gene matrix, the BS values for basal intergeneric nodes became weaker and the basal topology changed in each phylogeny as shown in supplementary Figures S8, S9 and S10.

### Tests for selection of chloroplast coding gene matrices

Tests performed on the chloroplast protein-coding genes were found to be under purifying/negative selection.

### Nuclear ITS cassette and ITS gene sequencing and alignment

The ITS cassette gene sequences of all six ingroup taxa and one outgroup were successfully generated from the ion semiconductor generated data. The ITS cassette region genes of *Bulnesia arborea* and gaps in the generated sequences of *Tribulus terrestris* and *Tetraena turbinata* ined. were successfully sequenced using Sanger sequencing. From these sequences an alignment matrix of ITS cassette gene sequences from six ingroup and two outgroup taxa was generated.

The amplification of the ITS region (ITS1-5SRNA-ITS2), Sanger sequencing, sequence assembly and alignment was successfully completed for the 59 taxa included in the study.

### Phylogenetic analyses of the nuclear rDNA gene sequences

The topologies of the phylogenetic trees of the nuclear rDNA ITS cassette region retrieved using maximum likelihood and Bayesian inference were identical but there were differences in clade support in the different analyses as shown in Figure S11. There is strong support for the monophyly of subfamily Zygophylloideae. The sister relationship of *Augea* and *Zygophyllum orbiculatum*/*stapffii* was strongly supported and this group was sister to *Roepera* with strong support. Relationships between *Zygophyllum, Fagonia* and *Tetraena* clades were not supported.

The phylogenetic tree of the nuclear rDNA ITS region is shown in Figure S12. Relationships within the genera are generally well supported. The resolution amongst genera is also much better than in the nuclear ITS cassette-based phylogeny.

Combining the ITS cassette and the ITS region data resulted in an overall well supported phylogeny (Figure 3) with the exception of the node between genus *Tetraena* and the clade consisting of the genera *Zygophyllum* and *Fagonia*.

**Figure 3:**
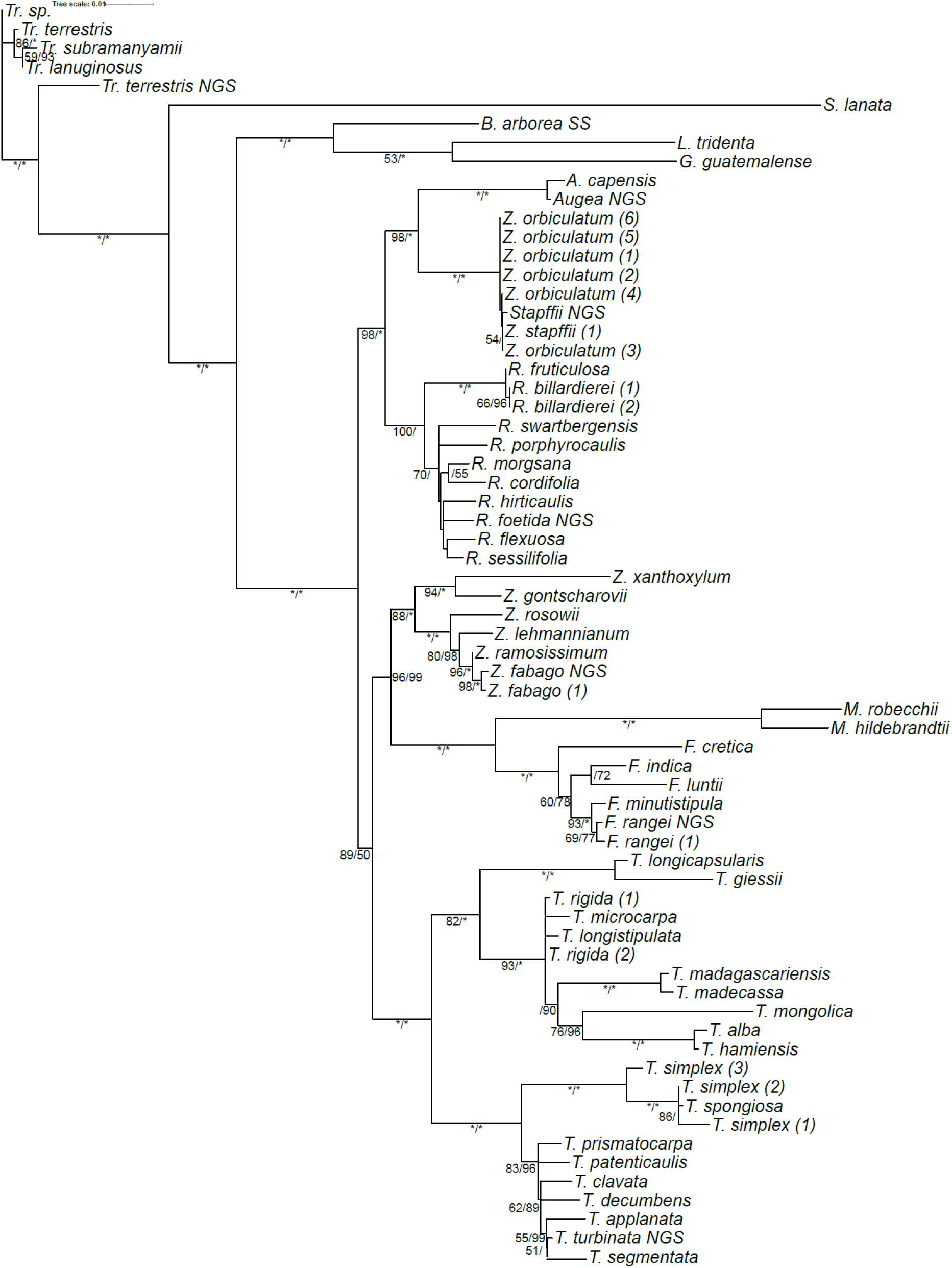
The most likely phylogenetic tree from the maximum likelihood analysis of the 67 taxa ITS cassette and ITS region combined alignment matrix. Maximum likelihood bootstrap values followed by Bayesian posterior probabilities are indicated on the nodes. Nodes with 100% bootstrap or posterior probabilities are indicated with an asterisk.

### Conflicts between phylogenies generated from chloroplast and nuclear gene matrices

The chloroplast and the nuclear gene tree topologies within the subfamily Zygophylloideae (Figures 3 and 5) showed conflict with regard the position of both the Asian *Zygophyllum* clade and the *Tetraena* clade. Consequently, this did not allow combined analysis of chloroplast and nuclear data.

### Molecular dating analysis

Since gene tree conflict precluded combined analysis, for the molecular clock analysis we opted to use the chloroplast sequence data matrix only. The dated phylogenetic tree generated from the 15 taxa 18NPG alignment matrix is shown in Figure 4. All of the nodes within the Zygophylloideae are strongly supported (PP>0.96). The age of the family Zygophylloideae is dated at 24.75 [15.06-36.71] Mya when *Fagonia* diverged from the rest of the subfamily. Subsequently, the genera *Zygophyllum*, *Tetraena* and the *Roepera*, *Zygophyllum orbiculatum*/*stapffii, Augea* group, radiated from the lineage by 20.76 [12.06-29.01] (basal node between *Tetraena* and the *Roepera*, *Zygophyllum orbiculatum*/s*tapffii, Augea* group) within the relatively short period of time of just under 4 million years indicating an ancient rapid radiation in the subfamily.

**Figure 4:**
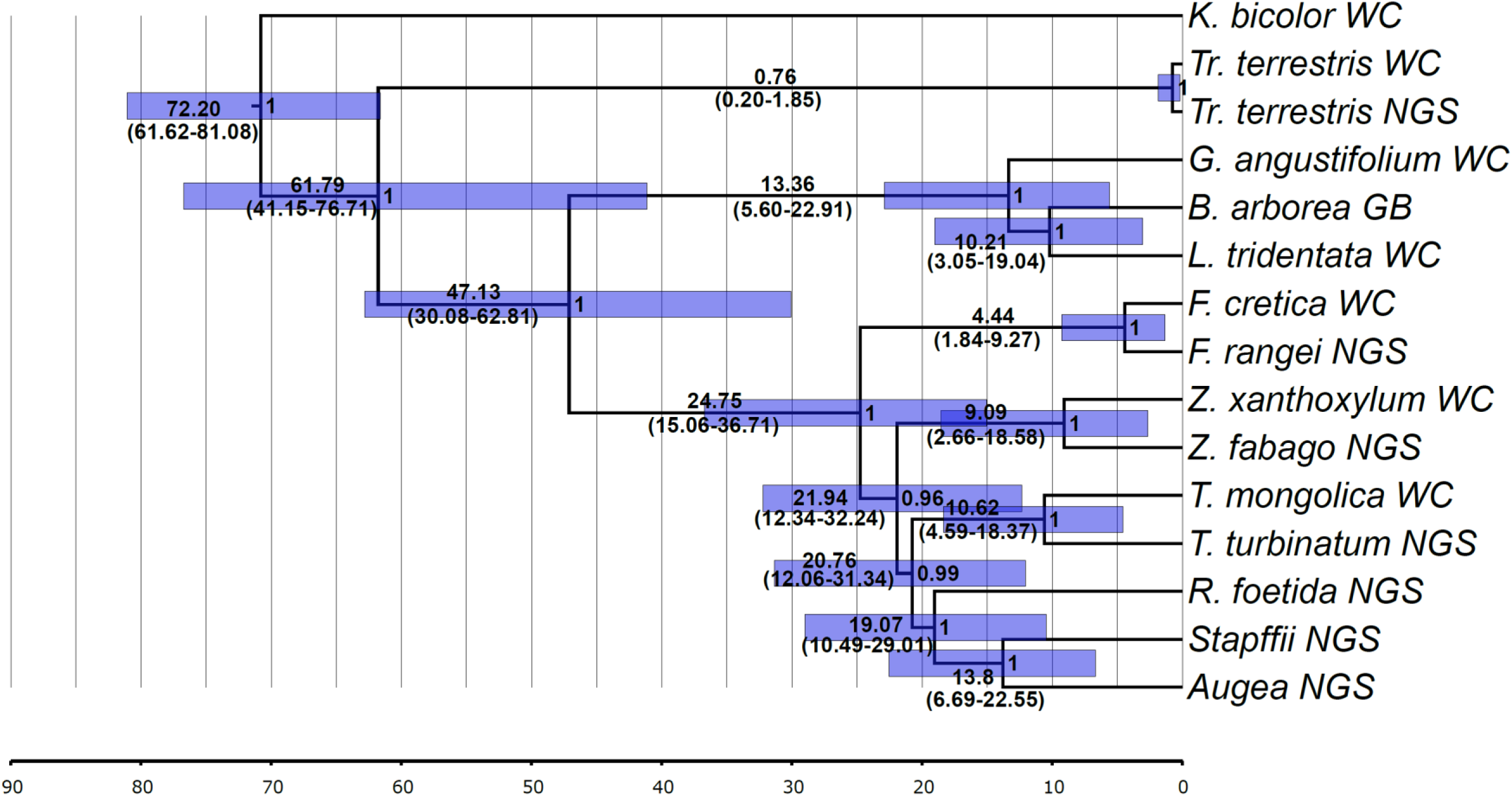
Maximum clade credibility tree inferred from BEAST based on the 15 taxa 18 coding genes alignment matrix to estimate the age of the Zygophylloideae radiation. Node support is indicated with posterior probability values, and the bars around node ages indicate 95% highest posterior density intervals.

## Discussion

What are the phylogenetic relationships between genera in the subfamily Zygophylloideae, and why have they been so difficult to infer? The lineage divergences have been dated to the early Miocene, around 20 million years ago, and likely occurred in quick succession (Bellstedt et al., 2012) which is challenging for phylogenetic inference. Previous studies using highly variable non-coding genes failed to resolve them (Bellstedt et al., 2008, Wu et al., 2015; Lauterbach et al., 2016; Wu et al., 2018) and although slowly evolving coding genes might offer more pertinent variation (Barrett et al., 2013, 2014, Zhang et al., 2016) prior to this work they had yet to be applied. Here, we tested both. Our results showed gene tree differences both between and apparently within genomes, which, in addition to a lack of informative characters, may explain the past uncertainty.

The chloroplast genome is generally inherited uniparentally, and in plant groups in which no recombination has been documented (including the subfamily Zygophylloideae), we might expect phylogenetic analysis of increasing numbers of chloroplast markers to give increasingly robust phylogenies (Givnish et al., 2018). Our analysis of 39 coding genes of 16 taxa representing all of the groups in the subfamily (C39, Figure 2), failed to resolve the outstanding uncertainties. By contrast, the phylogeny generated from the just 18 genes of the C18 matrix (excluding genes directly involved in the photosynthetic process; Figure 2 B) was almost completely supported. Why did less data result in a more robust hypothesis of relationships? Phylogenetic analysis of the other 21 coding genes (C21, Figure 2 C) directly involved in the photosynthetic process produced a phylogeny with discrete topological differences, but with conspicuously little support, particularly given the amount of data it represents. We did not identify any specific examples of convergence associated with shifts in the mode of photosynthesis (from C_3_ to C_4_ and to CAM) that might cause misleading phylogenetic signal. Tests for selection showed that these genes were under negative selection, and this is perhaps not unexpected in the absence of independent examples of C_4_ or CAM in the dataset. Given the otherwise unexpected negative impact of this subset of the data on phylogenetic inference, we can only speculate that some combination of functional constraints and selection acting on genes directly involved in the photosynthetic process hinders rather than helps us to infer the chloroplast phylogeny.

Lauterbach et al. (2016) showed that just three variable chloroplast gene alignments give a phylogeny with excellent resolution within genera of Zygophylloideae, but little intergeneric resolution (Figure S2 here generated from a re-aligned sequence matrix). Addition of the sequences of nine noncoding gene regions (16 taxa 5NC and 3HVNC) to the 16 taxa 18NPG gene matrix followed by phylogenetic analysis also gives a largely resolved phylogeny (Figure S1), but with inconsistent support for two apparently conflicting but very short internal branches (Figure S1 vs Figure 2 B). Addition of even greater numbers of 3HVNC genes to the 5NC genes of 16 taxa does not change the topology nor increase support values of the spine of the phylogeny (supplementary Figure S3, S4 and S5), neither does adding more such data from additional taxa (34 taxa 3HVNC) to the 16 taxa 39C + 5NC + 3HVNC (Figure S6) phylogeny (64 taxa, Figure S7, and 122 taxa, Figure S8). Despite the potential with denser taxon sampling to improve alignments and reveal informative sites, these regions also appear uninformative at this level.

Perhaps nuclear gene sequences may be better suited for phylogenetic inference in the subfamily? Our ITS cassette region data on its own delivers a phylogeny with less support than the 18NPG chloroplast phylogeny but is based on far less sequence data (compare alignment lengths in Table 1) and, perhaps less than ideally, fewer taxa (8 taxa as opposed to the 16 used to establish the chloroplast phylogenies). Addition of noncoding ITS sequences to coding (rRNA) gene sequences fails to resolve all the basal nodes. However, the addition of large numbers of taxa with noncoding genes to a smaller number of taxa with coding genes resulted in a large, combined phylogeny that is better supported (Figure 3) than that based on ITS sequences alone. This is in contrast to what we have found for the chloroplast genome when we combined the sequence data of many coding genes from a limited number of taxa with non-coding gene sequences of a much larger number of taxa (i.e. that this weakened rather than strengthened intergeneric resolution).

Better resolved chloroplast (Figure 2 B) and nuclear (Figure 3) phylogenies of Zygophylloideae revealed supported topological differences. This could reflect either ancient introgressive hybridization or incomplete lineage sorting. We have not attempted to test which of these mechanisms could be responsible: the data represent just two independent gene trees (precluding e.g. ABBA BABA frequency tests; e.g. Yardeni et al., 2025), and coalescent simulation approaches would depend on assumptions about population sizes through a long time period and would be unlikely to reject coalescence (De Villiers et al., 2013). With datasets representing few independent gene trees, causes of gene tree conflict may remain uncertain (e.g., Barrett et al., 2013; Bell et al., 2010; Sun et al., 2015, Xi et al., 2014). Recently, the Angiosperms353 project has included taxa from the Zygophylloideae in their analyses (Baker et al., 2022) and the results obtained with a similar choice of taxa yield a phylogeny of which the topology is identical to that of the chloroplast 18NPG topology obtained in this study, as opposed to either the (unsupported) plastid photosynthesis gene topology or nuclear ITS based phylogeny. This could be interpreted as independent support for the 18NPG chloroplast phylogenetic hypothesis best representing the underlying species tree (which also happens to be the topology represented in our molecular dating analyses). However, it would not exclude the possibility of processes such as ancient introgressive hybridization or incomplete lineage sorting in explaining individual gene tree differences.

The short internal branch lengths between different clades in the Zygophylloideae subtending long stem lineages is indicative of a rapid ancient radiation. Divergence between the genera in the subfamily Zygophylloideae occurred in quick succession after 24.75 Mya (Figure 4). This coincides with the start of the worldwide Miocene aridification (Zachos et al., 2001). Some of these lineages radiated into larger genera (*Fagonia, Tetraena*, *Zygophyllum* and *Roepera*) occupying a variety of habitats and habitat niches (Lauterbach et al., 2016). The clade of monotypic *Augea*, *Zygophyllum orbiculatum/stapffii* and *Roepera,* supported in both the nuclear and chloroplast phylogenies (when excluding photosynthesis genes), represent a southern African radiation. Two of these lineages are presently represented by single species with widespread distributions (i.e. *Augea capensis* and *Zygophyllum orbiculatum/stapffii*). In other Zygophyllaceae subfamilies, in the Tribuloideae and Seetzeniodieae, there is a similar pattern of lineages that have radiated widely across arid biomes, as in *Tribulus* (Lauterbach et al., 2019), and the monotypic genera *Sisyndite spartea* E.Mey. ex Sond. and *Neoluederitzia sericeocarpa* Schinz.

The species with the widest geographical distribution in the Zygophylloideae (from southern- to northern Africa including the Azores) is *Tetraena simplex* (Sheahan & Chase, 2000). It is an annual prostrate herb with a tumbleweed seed distribution mechanism and C_4_ photosynthesis*. Roepera cordifolia* that exhibits CAM photosynthesis (Matimati et al. 2012) also has a large range and has dispersed into the hyper-arid southern Namib desert in Namibia. *Tribulus terrestris*, another example of C_4_, in the Tribuloideae, is distributed almost worldwide (Lauterbach et al., 2019). In ancestral lineages, preadaptation to aridity potentially including arid adapted photosynthetic mechanisms may have allowed the Zygophylloideae to disperse from the arid areas in southern Africa into similar arid areas in northern Africa and Asia via the African Arid Corridor, to Madagascar, and more recently into Australia by long distance dispersal (Bellstedt et al., 2012). Sequence variation that reflects shifts in photosynthetic mode may therefore be directly linked to important phenomena such as drought tolerance and potential for dispersal rather than accurately reflecting their phylogenetic relationships.

## Conclusion

Phylogenetic relationships within the subfamily Zygophylloideae have been a point of contention for many years, as neither multiple linked chloroplast genes nor single nuclear marker molecular studies proved conclusive. Our results go some way to explain why: First, the lack of consistent congruent phylogenetic signal in photosynthesis related genes compared to the rest of the chloroplast-encoded markers. This will have impacted previous studies that relied on *rbc*L (e.g., Bellstedt et al., 2012) and apparently can be problematic even in the context of much larger datasets such as ours. Second, nuclear-chloroplast conflict, as also reported e.g. by Wu et al. (2015) and in subfamily Tribuloideae by Lauterbach et al. (2019), which suggests, paraphrasing Hahn and Nakhleh (2016), that we should temper our ‘exuberance’ for inferring a meaningful bifurcating species tree.

The data analysed here goes well beyond the few Sanger sequenced markers used in most previous studies of Zygophylloideae, and the subset of chloroplast genes not related to photosynthesis provided robust support for a topology that is consistent with independent data (Angiosperm353: Baker et al., 2022). Despite conflict, this might be useful as a hypothesis for a species tree.

Nevertheless, our results represent just two independent gene trees and (for most of the sequence data) minimal taxon sampling, and we were unable to pinpoint the phenomena underlying the gene tree conflict. We need to better represent the species diversity of Zygophylloideae in analyses of multiple independent gene trees to address this and further improve our understanding of relationships, whether they are tree-like, or not. Given the results presented here, we would urge the careful interpretation of both gene tree conflict and gene function in such analyses.

## Acknowledgements

Dr Felix Forest, the Royal Botanic Gardens, Kew and Dr Michael Moeller, Royal Botanic Garden Edinburgh are thanked for the plant and DNA samples they provided. Cape Nature, Northern Cape Department of Nature Conservation and Environmental Conservation, and the Namibian Ministry of Environment and Tourism are thanked for permission to collect material. A preprint version of this article has been peer-reviewed and recommended by PCI Evol Biol (https://doi.org/10.24072/pci.evolbiol.101119). We thank the Recommender, Beniot Nabholz, the reviewer, Sidonie Bellot, and two anonymous reviewers for very helpful comments on previous versions of this paper.

## Data, scripts, code, and supplementary information availability

Raw data are available online at NCBI under the bioproject, PRJNA1526063 (https://www.ncbi.nlm.nih.gov/bioproject/?term=prjna1526063). All *de novo* assembled contiguous chloroplast, nuclear and mitochondrial sequences and raw reads were deposited in the Stellenbosch University data repository (https://doi.org/10.25413/sun.31238719). All supplementary files and tree files are available on Zenodo (https://doi.org/10.5281/zenodo.23560646).

## Supplementary Tables

**Supplementary Table 1:** Collection details of species used for ion semi-conductor sequencing.

| **Species** | **Collection locality** | **Country** | **Collectors** |
| --- | --- | --- | --- |
| *Augea capensis* | Tankwa Karoo near Ceres, Western Cape, South Africa. (Bellstedt, 934) | South Africa | Dirk U. Bellstedt & P. De Wet van der Merwe |
| *Fagonia rangei* | Close to embankment of the Orange River near Vioolsdrif border outpost between South Africa and Namibia. (Namibian side) (Bellstedt, 1314) | Namibia | Dirk U. Bellstedt |
| *Roepera foetida* | Bridge across the Breede River between Robertson and McGregor Western Cape, South Africa (Robertson side) (Bellstedt 1387) | South Africa | P. De Wet van der Merwe |
| *Tetraena turbinata* ined. | Tankwa Karoo near Ceres, Western Cape, South Africa. (Bellstedt, 1386) | South Africa | Dirk U. Bellstedt & P. De Wet van der Merwe |
| *Tribulus terrestris* | Stellenbosch Train station, Western Cape, South Africa. (P.D.W. van der Merwe, s.n.) | South Africa | Dirk U. Bellstedt & P. De Wet van der Merwe |
| *Zygophyllum fabago* | Germany (Botanical Garden, Mainz University), Native to the Middle East. | Germany | Michael D. Pirie |
| *Zygophyllum stapffii* | Near Swakopmund, Namibia. (Bellstedt, 1222) | Namibia | Dirk U. Bellstedt |

**Supplementary Table 2:**
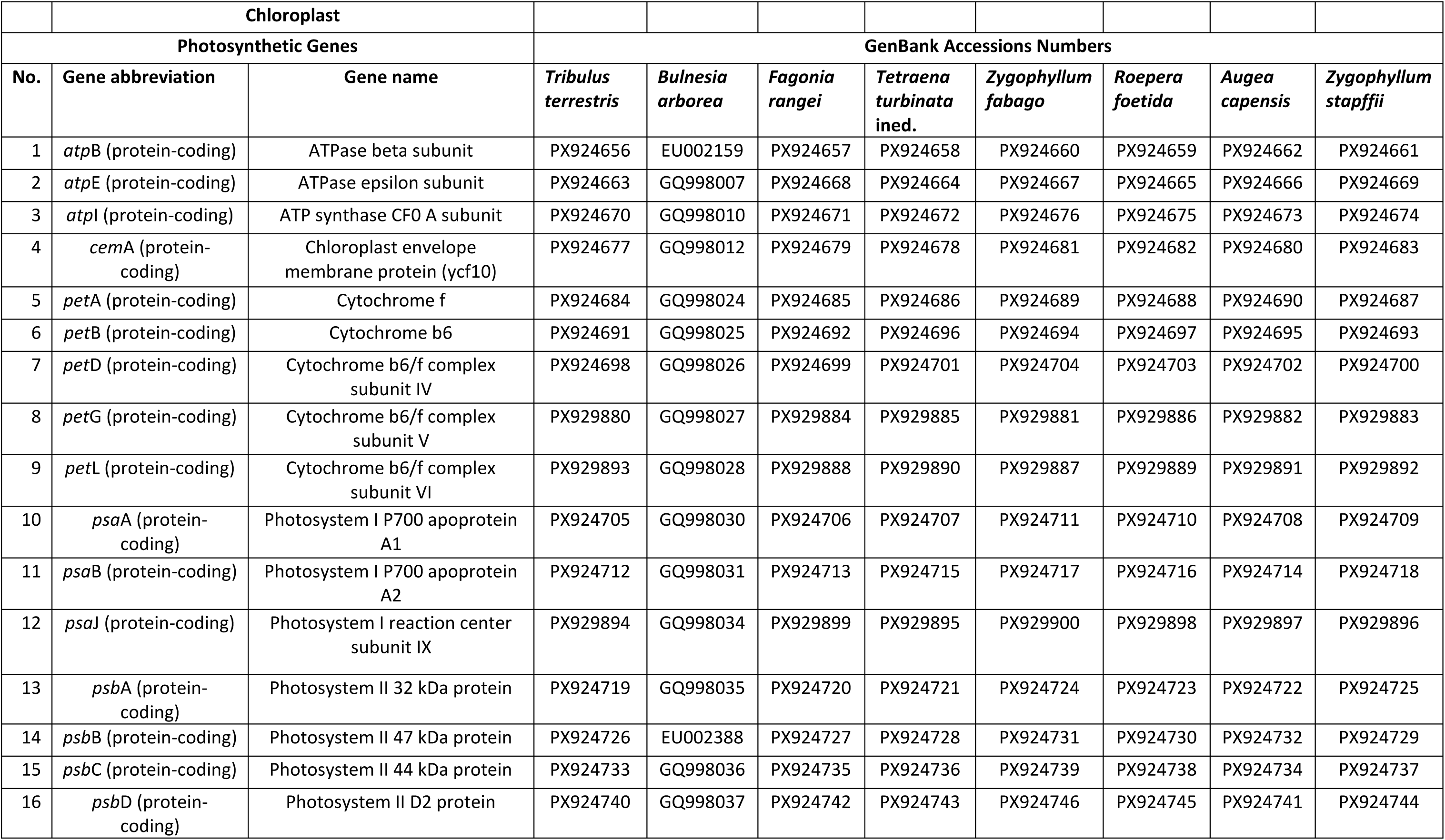
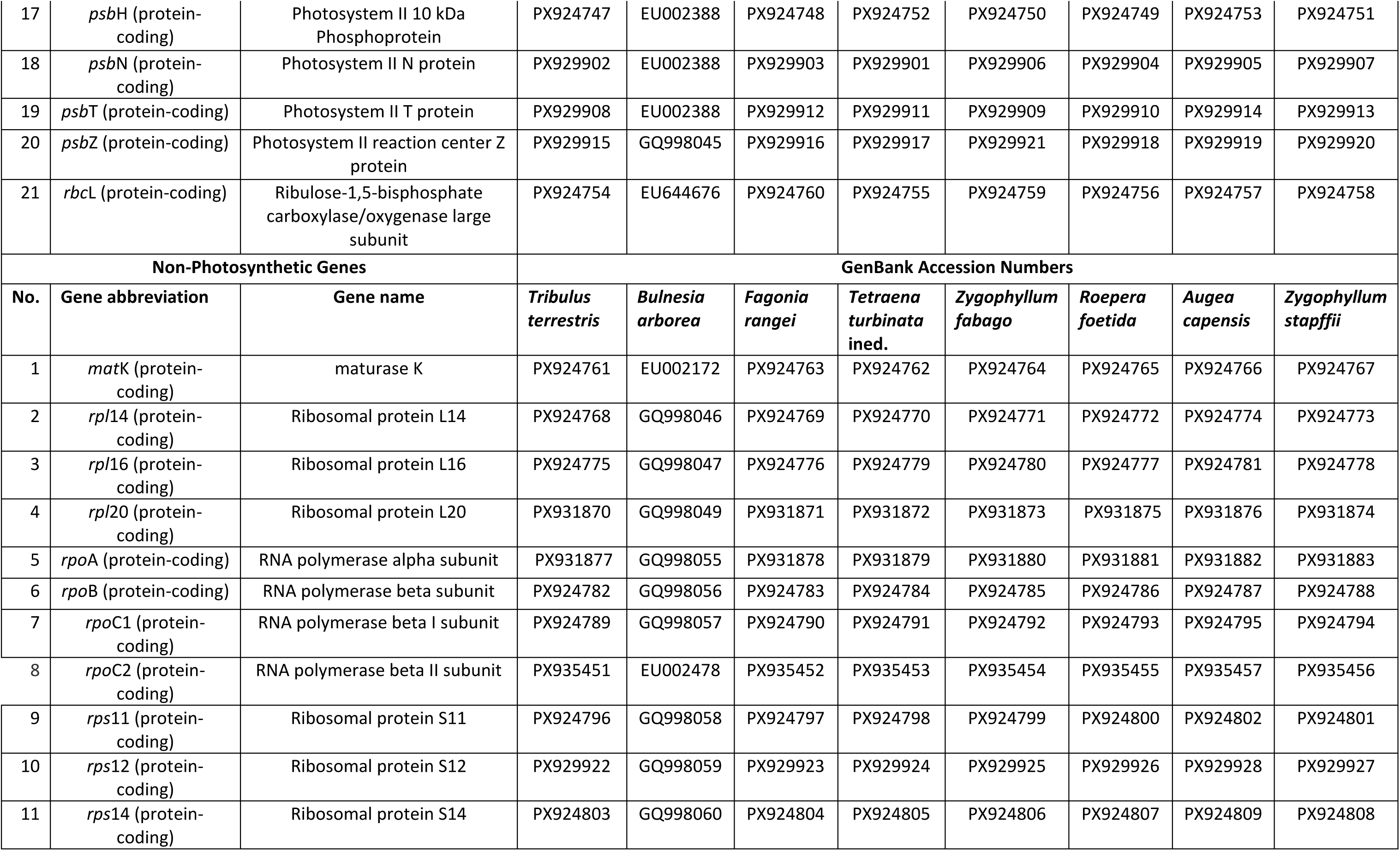
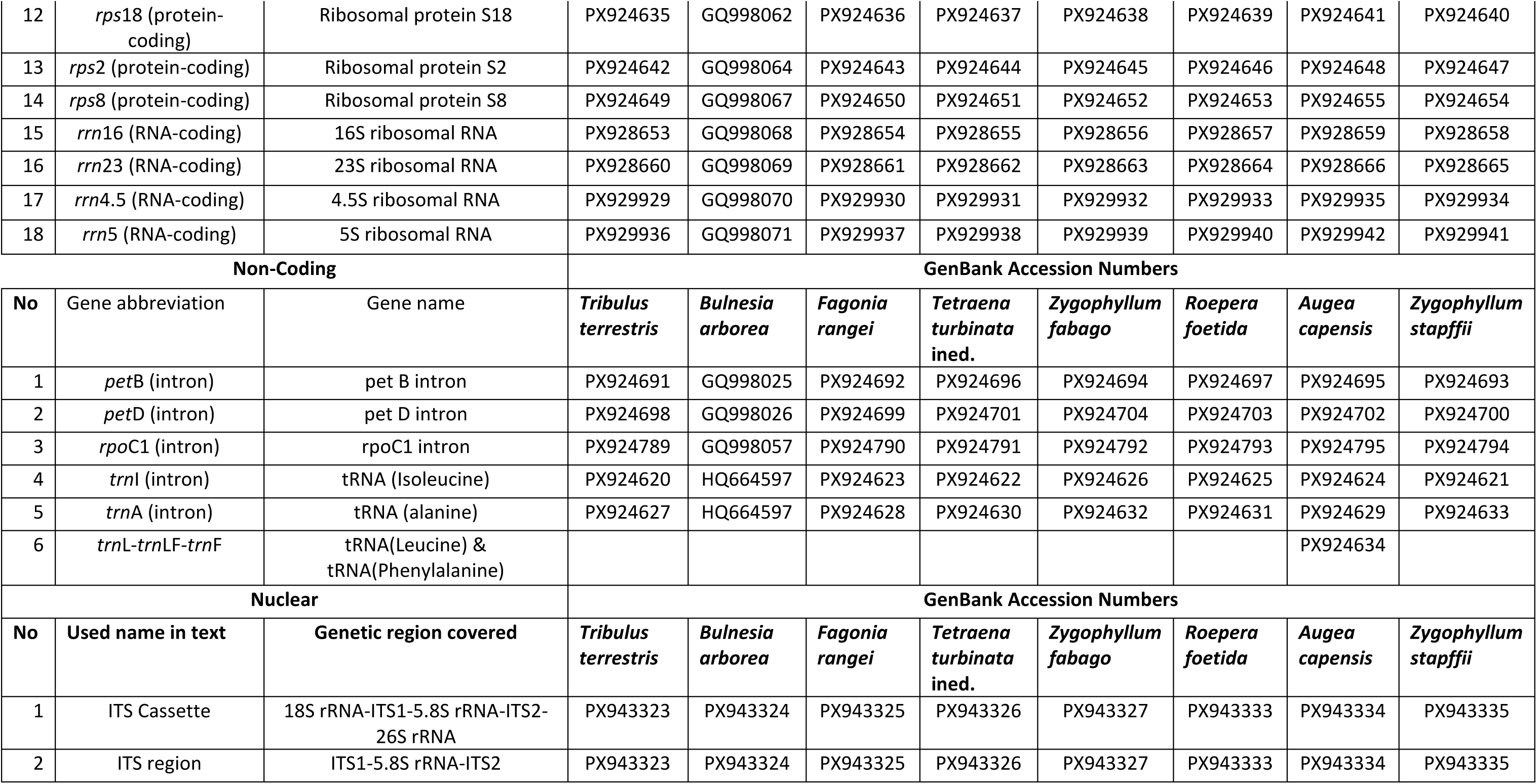
GenBank accession codes of gene sequences used.

**Supplementary Table 3:** Collection details of all species used to generate ITS sequences in this study and their ITS GenBank accession numbers.

| No. | Species | Name on trees | Geographic distribution | Voucher details | Collection locality | Accession number |
| --- | --- | --- | --- | --- | --- | --- |
| 1 | *Augea capensis* | *A. capensis* | South Africa | Bellstedt (STE) | Calitzdorp, South Africa | PX943336 |
| 2 | *Augea capensis* | *Augea* NGS | South Africa | Bellstedt 934 | Tankwa Karoo near Ceres, Western Cape | PX943334 |
| 3 | *Bulnesia arborea* | *B. arborea* SS | New World | Michael Moeller | Royal Botanical Gardens, Edinburgh | PX943324 |
| 4 | *Fagonia cretica* | *F. cretica* | Canary Islands | Beier 125 (UPS) |  | AY641623 |
| 5 | *Fagonia indica* | *F. indica* | United Arab Emirates | Thulin et al. 10024 (UPS) |  | AY641631 |
| 6 | *Fagonia luntii* | *F. luntii* | Yemen | Thulin et al. 9881 (UPS) |  | AY641638 |
| 7 | *Fagonia minutistipula* | *F. minutistipula* | Namibia | Giess and Muller 13952 |  | AY641641 |
| 8 | *Fagonia rangei* | *F. rangei* (1) | South Africa | Leistner 3388 |  | AY641647 |
| 9 | *Fagonia rangei* | *F. rangei* NGS | Namibia | Bellstedt 1314 | Close to embankment of the Orange River near Vioolsdrif border outpost between South Africa and Namibia, South Africa | PX943325 |
| 10 | *Guaiacum guatemalense* | *G. guatemalense* | New World | Chase 640 (K) |  | PX943337 |
| 11 | *Larrea tridentata* | *L. tridentata* | New World | Chase 636 (K) |  | PX943338 |
| 12 | *Melocarpum hildebrandtii* | *M. hildebrandtii* | Somalia | Thulin et al. 9012 (UPS) |  | AY641615 |
| 13 | *Melocarpum robecchii* | *M. robecchii* | Yemen | Thulin et al. 9537 (UPS) |  | AY641616 |
| 14 | *Roepera billardierei* (1) | *R. billardierei* (1) | Australia | SR.417 (Adelaide BG.) | Adelaide | AY641613 |
| 15 | *Roepera billardierei* (2) | *R. billardierei* (2) | Australia | 2287 |  | PX943344 |
| 16 | *Roepera cordifolia* | *R. cordifolia* | South Africa | Marais 446 (STE) | Koekenaap, South Africa | PX943346 |
| 17 | *Roepera flexuosa* | *R. flexuosa* | South Africa | Bellstedt 794 (STE) | Langebaan, WC, South Africa | PX943348 |
| 18 | *Roepera foetida* | *R. foetida* NGS | South Africa | Bellstedt 1387 | Bridge across the Breede River between Robertson and McGregor Western Cape, (Robertson side), South Africa | PX943333 |
| 19 | *Roepera fruticulosa* | *R. fruticulosa* | W. Australia | Chase 2203 (K) | Port Gregory, North of Perth | PX943349 |
| 20 | *Roepera hirticaulis* | *R. hirticaulis* | Namibia | Craven 2857 |  | PX943352 |
| 21 | *Roepera margsana* | *R. margsana* | South Africa | Bellstedt 890 (STE) | Steinkopf, NC, South Africa | PX943358 |
| 22 | *Roepera porphyrocaulis* | *R. porphyrocaulis* | South Africa | Bellstedt 800 (STE) | Inland WC | PX943366 |
| 23 | *Roepera sessilifolia* | *R. sessilifolia* | South Africa | Marais 434 (STE) | Morreesburg, WC, South Africa | PX943372 |
| 24 | *Roepera swartbergensis* | *R. swartbergensis* | South Africa | Bellstedt 798 (STE) | Swartberg Pass, WC, South Africa | PX943378 |
| 25 | *Seetzenia lanata* | *S. lanata* | South Africa | Bellstedt 938 | Koue Bokkeveld, Western Cape, South Africa | PX943339 |
| 26 | *Tetraena alba* | *T. alba* | Yemen | Thulin et al. 7977 (1697) (K) |  | PX943342 |
| 27 | *Tetraena applanata* | *T. applanata* | Namibia | Bellstedt 870 (STE) | Rosh Pinah, Namibia | PX943343 |
| 28 | *Tetraena clavata* | *T. clavata* | Namibia | Bellstedt 878 (STE) | Lüderitz peninsula, Namibia | PX943345 |
| 29 | *Tetraena decumbens* | *T. decumbens* | Namibia | Bellstedt 851 (STE) | Namibia | PX943347 |
| 30 | *Tetraena giessii* | *T. giessii* | Namibia | Bellstedt 1326 | Arimas Farm, Witputz, Namibia | PX943350 |
| 31 | *Tetraena hamiensis* | *T. hamiensis* | Yemen | Thulin and Berer 9840 (UPS) |  | PX943351 |
| 32 | *Tetraena longicapsularis* | *T. longicapsularis* | Namibia | Bellstedt 879 (STE) | Lüderitz peninsula. Namibia | PX943353 |
| 33 | *Tetraena longistipulata* | *T. longistipulata* | Namibia | Bellstedt 1322 | Arimas Farm, Witputz, Namibia | PX943354 |
| 34 | *Tetraena madagascariensis* | *T. madagascariensis* | Madagascar | Bellstedt 1239 | Tulear district, Madagascar | PX943355 |
| 35 | *Tetraena madecassa* | *T. madecassa* | Madagascar | Bellstedt 1243 | Taolagnaro District, Madagascar | PX943356 |
| 36 | *Tetraena microcarpa* | *T. microcarpa* | Namibia | Bellstedt 1315 | Arimas Farm, Witputz, Namibia | PX943357 |
| 37 | *Tetraena mongolica* | *T. mongolica* | Mongolia | Sheahan 1105 | Dung Kou, Inner Mongolia | PX943340 |
| 38 | *Tetraena patenticaulis* | *T. patenticaulis* | Namibia | Bellstedt 868 (STE) | Rosh Pinah, Namibia | PX943365 |
| 39 | *Tetraena prismatocarpa* | *T. prismatocarpa* | Namibia | Bellstedt 863 (STE) | Dumusibriver, near Orangeriver, Namibia | PX943367 |
| 40 | *Tetraena rigidum* | *T. rigidum* (1) | Namibia | Bellstedt 852 (STE) | Noordoewer, Namibia | PX943368 |
| 41 | *Tetraena rigidum* | *T. rigidum* (2) | Namibia | Bellstedt 855 (STE) | Aussenkehr, Namibia | PX943369 |
| 42 | *Tetraena segmentata* | *T. segmentata* | Namibia | Bellstedt 861 (STE) | Rosh Pinah, Namibia | PX943371 |
| 43 | *Tetraena simplex* | *T. simplex* (1) | Namibia | Bellstedt 1226 (STE) | Grünau, Namibia | PX943374 |
| 44 | *Tetraena simplex* | *T. simplex* (2) | Namibia | Bellstedt 1224 (STE) | Swakoprivier mouth, Namibia | PX943373 |
| 45 | *Tetraena simplex* | *T. simplex* (3) | Africa and Asia | El Hadidi, Sheahan 806 | Northern Hemisphere | PX943375 |
| 46 | *Tetraena simplex* (*spongiosa*) | *T. spongiosa* | Angola | Bellstedt 1214 (STE) | Praia Azul, Benguela, Angola | PX943376 |
| 47 | *Tetraena turbinata* ined. | *T. turbinata* NGS | South Africa | Bellstedt 1386 | Tankwa Karoo near Ceres, Western Cape | PX943326 |
| 48 | *Tribulus lanuginosus* | *Tr. lanuginosus* |  | Sathishkumar,R, et al. |  | HM236860 |
| 49 | *Tribulus* sp. | *Tr. sp.* | Namibia | Bellstedt 1333 |  | PX943341 |
| 50 | *Tribulus subramanyamii* | *Tr. subramanyamii* |  | Sathishkumar,R, et al. |  | HM236858 |
| 51 | *Tribulus terrestris* | *Tr. terrestris* (1) |  | B. B. Simpson 16-VI-00-1 | USA: TX, Burnet Co | AY260972 |
| 52 | *Tribulus terrestris* | *Tribulus terrestris* NGS | South Africa |  | Stellenbosch Train station, Western Cape, South Africa. (P.D.W. van der Merwe, s.n.) | PX943323 |
| 53 | *Zygophyllum fabago* | *Z. fabago* (1) | France | Chase 516 (K) | Lyons Bot. Gard. | PX943328 |
| 54 | *Zygophyllum fabago* | *Z. fabago* NGS | Germany (Native to the Middle East) | MP | Botanical Garden, Mainz University) | PX943327 |
| 55 | *Zygophyllum gontscharovii* | *Z. gontscharovii* | Tajikistan | Ashnova 10191 (K) |  | PX943330 |
| 56 | *Zygophyllum lehmannianum* | *Z. lehmannianum* | Turkmenistan | 10192 (K) |  | PX943332 |
| 57 | *Zygophyllum orbiculatum* | *Z. orbiculatum* (1) | S. Angola | Bellstedt 1202 (STE) | Namibe District, Angola | PX943359 |
| 58 | *Zygophyllum orbiculatum* | *Z. orbiculatum* (5) | S. Angola | Craven 5096 (WIND) | 30 km west of Foz de Cunene, Angola | PX943363 |
| 59 | *Zygophyllum orbiculatum* | *Z. orbiculatum* (6) | Angola | Craven 5101 (WIND) | River tributary, South of Tambor, Angola | PX943364 |
| 60 | *Zygophyllum orbiculatum* | *Z. orbiculatum* (2) | S. Angola | Bellstedt 1213 (STE) | Lake Acoa, Angola | PX943360 |
| 61 | *Zygophyllum orbiculatum* | *Z. orbiculatum* (4) | Namibia | HK 2757 (WIND) |  | PX943362 |
| 62 | *Zygophyllum orbiculatum* | *Z. orbiculatum* (3) | Namibia | HK 2753 (WIND) |  | PX943361 |
| 63 | *Zygophyllum ramosissimum* | *Z. ramosissimum* | Kazakhstan | Giamolov 10182 (K) |  | PX943331 |
| 64 | *Zygophyllum rosowii* | *Z. rosowii* | China | Chaney 57 | Karakum | PX943370 |
| 65 | *Zygophyllum stapffii* | *Z. stapffii* | Namibia | Bellstedt 1222 (STE) | Swakopmund, Namibia | PX943377 |
| 66 | *Zygophyllum stapffii* | *Stapffii* NGS | Namibia | Bellstedt 1222 | Near Swakopmund, Namibia | PX943335 |
| 67 | *Zygophyllum xanthoxylum* | *Z. xanthoxylum* | China | Chase 1700 (K) |  | PX943329 |

**Supplementary Table 4:**
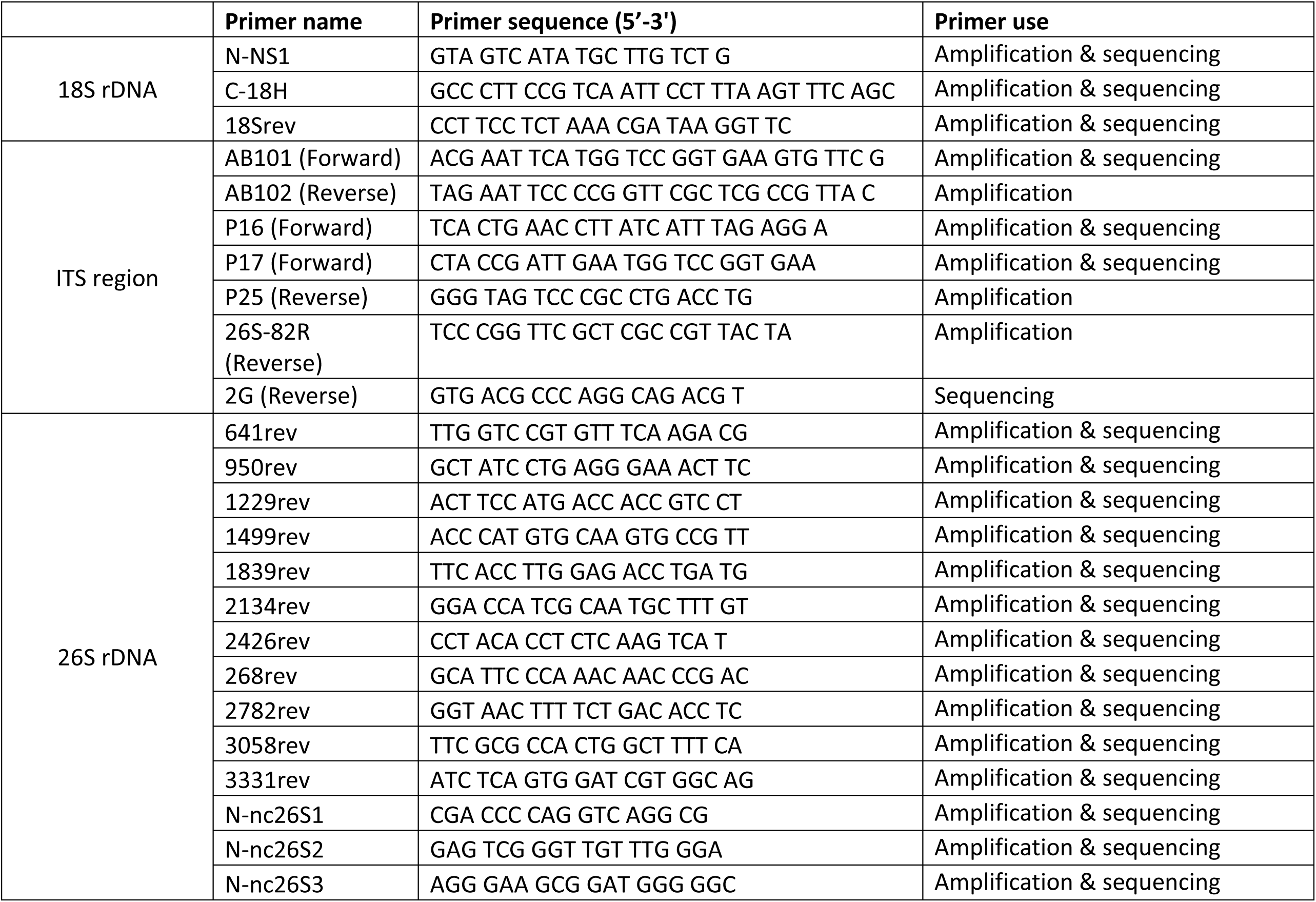
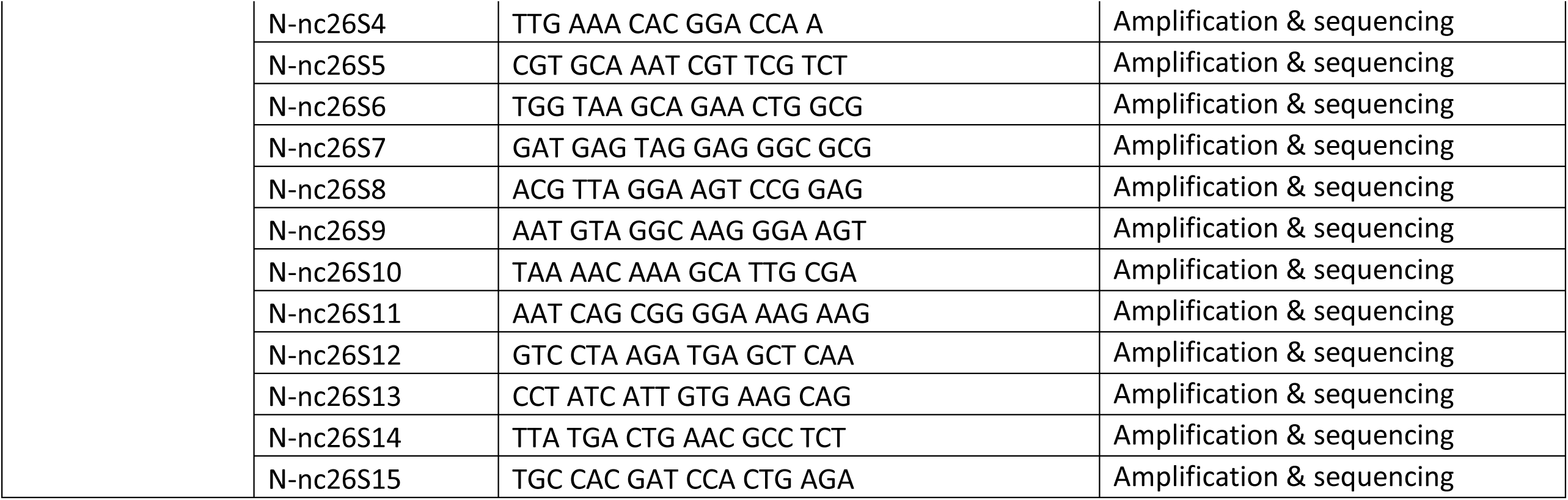
Primers used for sequencing ITS genes in *Bulnesia arborea*.

## Supplementary Figures

**Figure S1:**
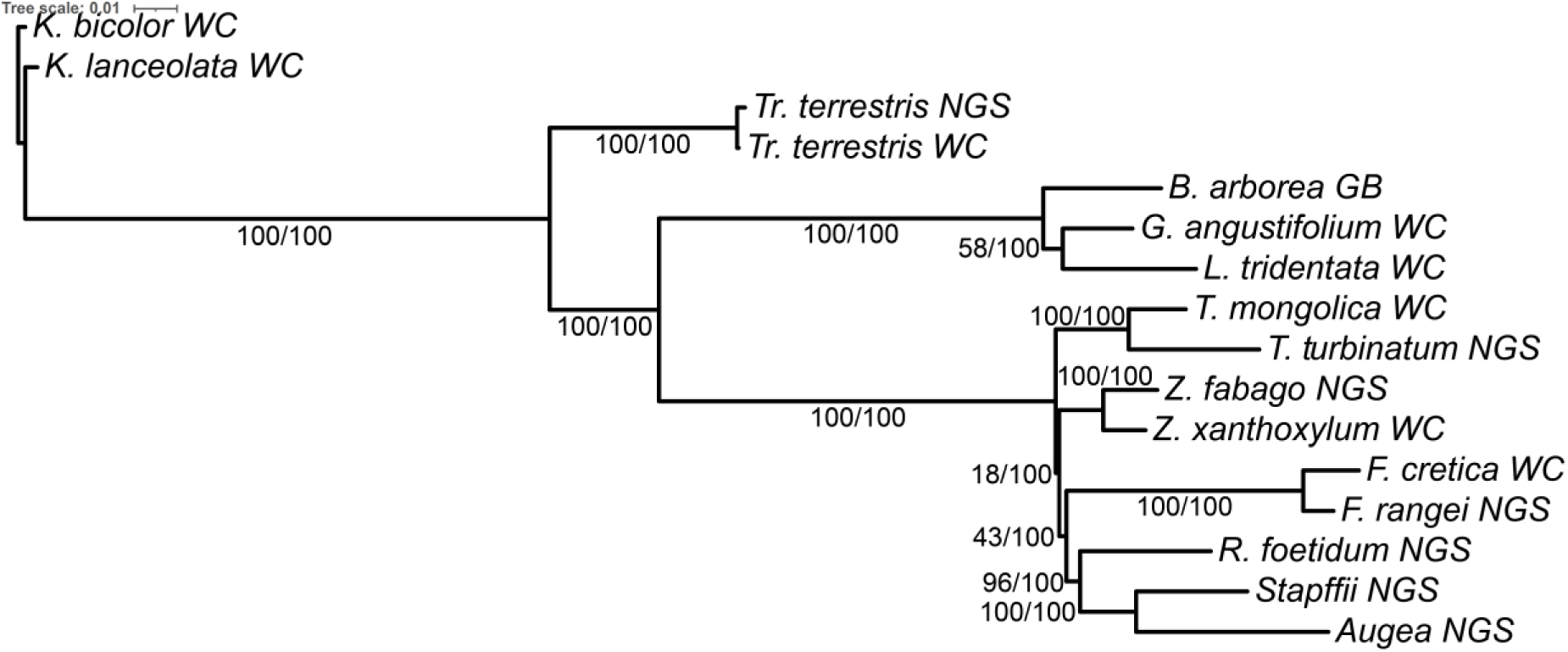
The most likely phylogenetic tree from the maximum likelihood analysis of the 16 taxa 5 NC gene and 3 HVNC gene alignment matrix. Maximum likelihood bootstrap values followed by Bayesian posterior probabilities are indicated on the nodes.

**Figure S2:**
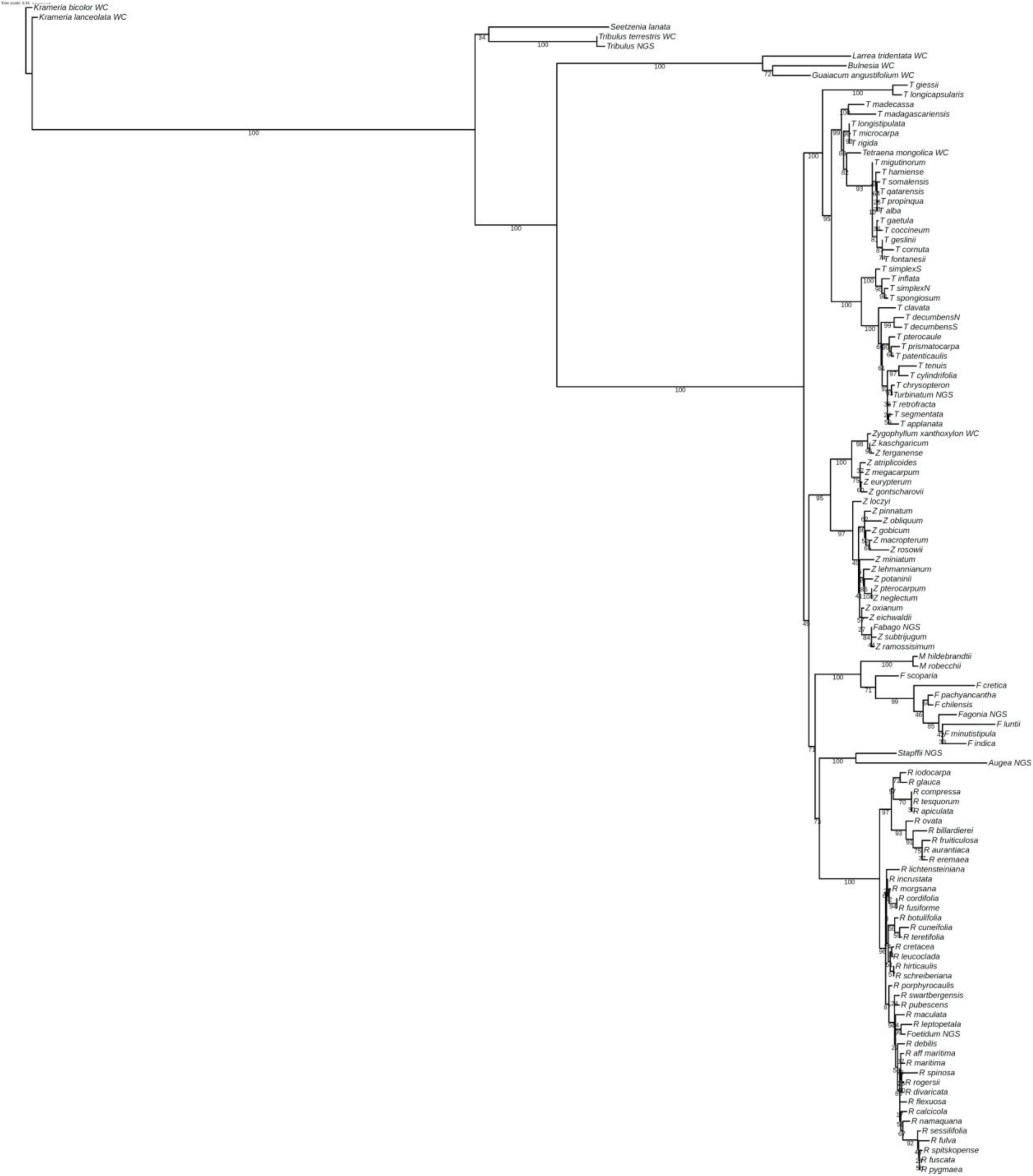
The most likely phylogenetic tree from the maximum likelihood analysis of the 3 HVNC gene alignment matrix of 9 ingroup taxa and 8 outgroups plus 105 species = 121 taxa. Maximum likelihood bootstrap values are indicated on the nodes.

**Figure S3:**
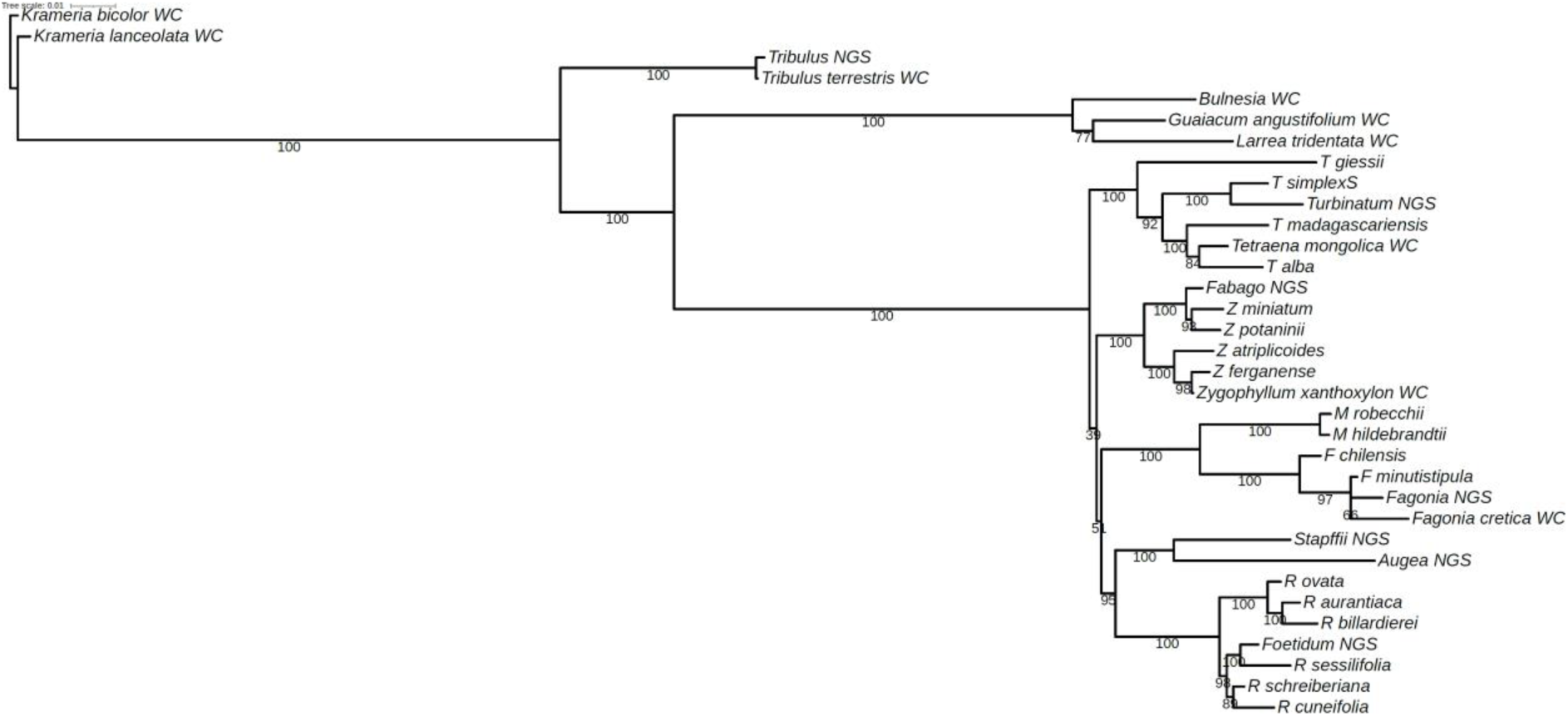
The most likely phylogenetic tree from the maximum likelihood analysis of the 3 HVNC genes plus 5 NC gene alignment matrix of 9 ingroup taxa and 7 outgroups plus 18 species = 34 taxa. Maximum likelihood bootstrap values are indicated on the nodes.

**Figure S4:**
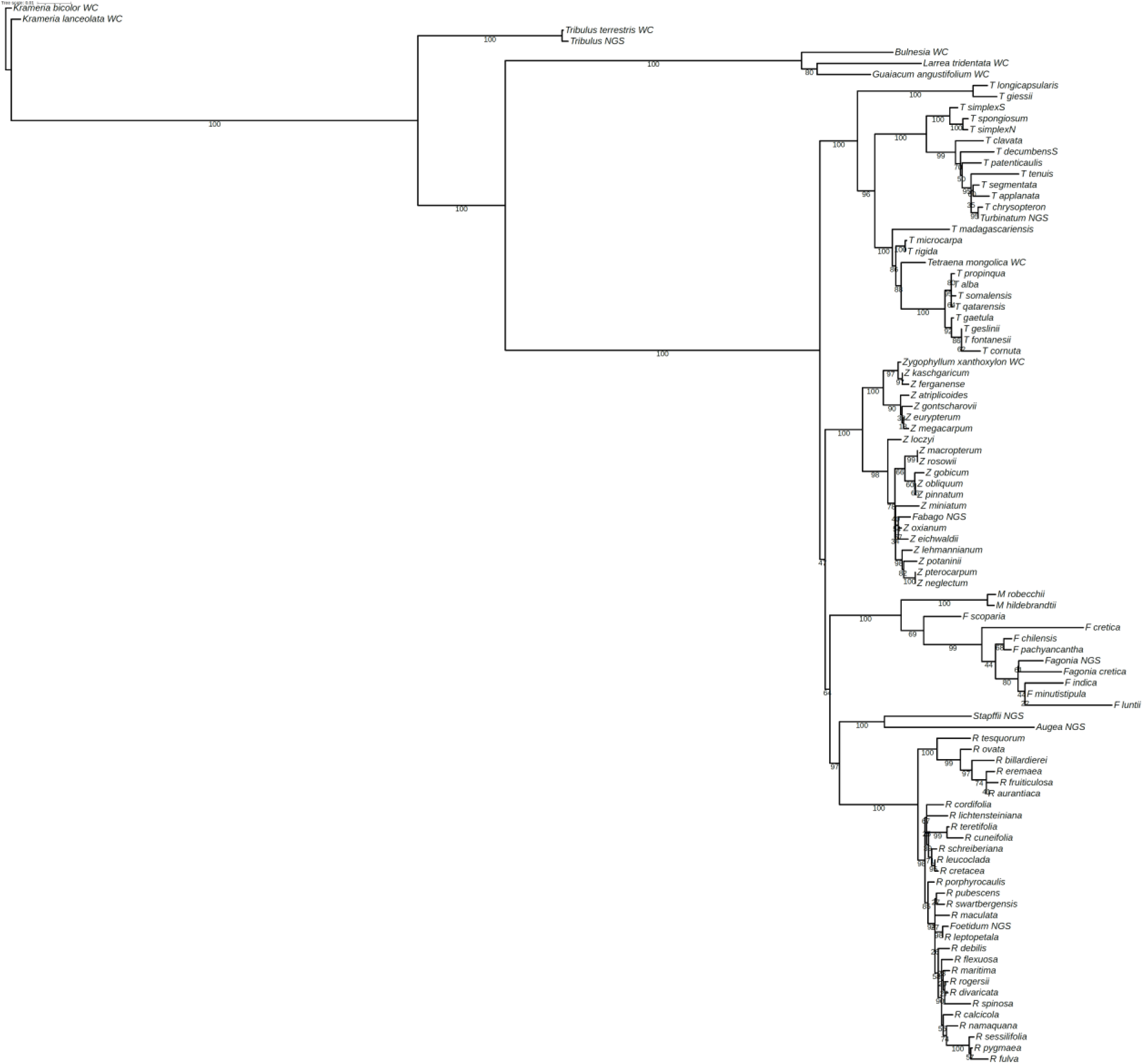
The most likely phylogenetic tree from the maximum likelihood analysis of the 3 HVNC genes plus 5 NC gene alignment matrix of 9 ingroup taxa and 7 outgroups plus 80 species = 96 taxa. Maximum likelihood bootstrap values are indicated on the nodes.

**Figure S5:**
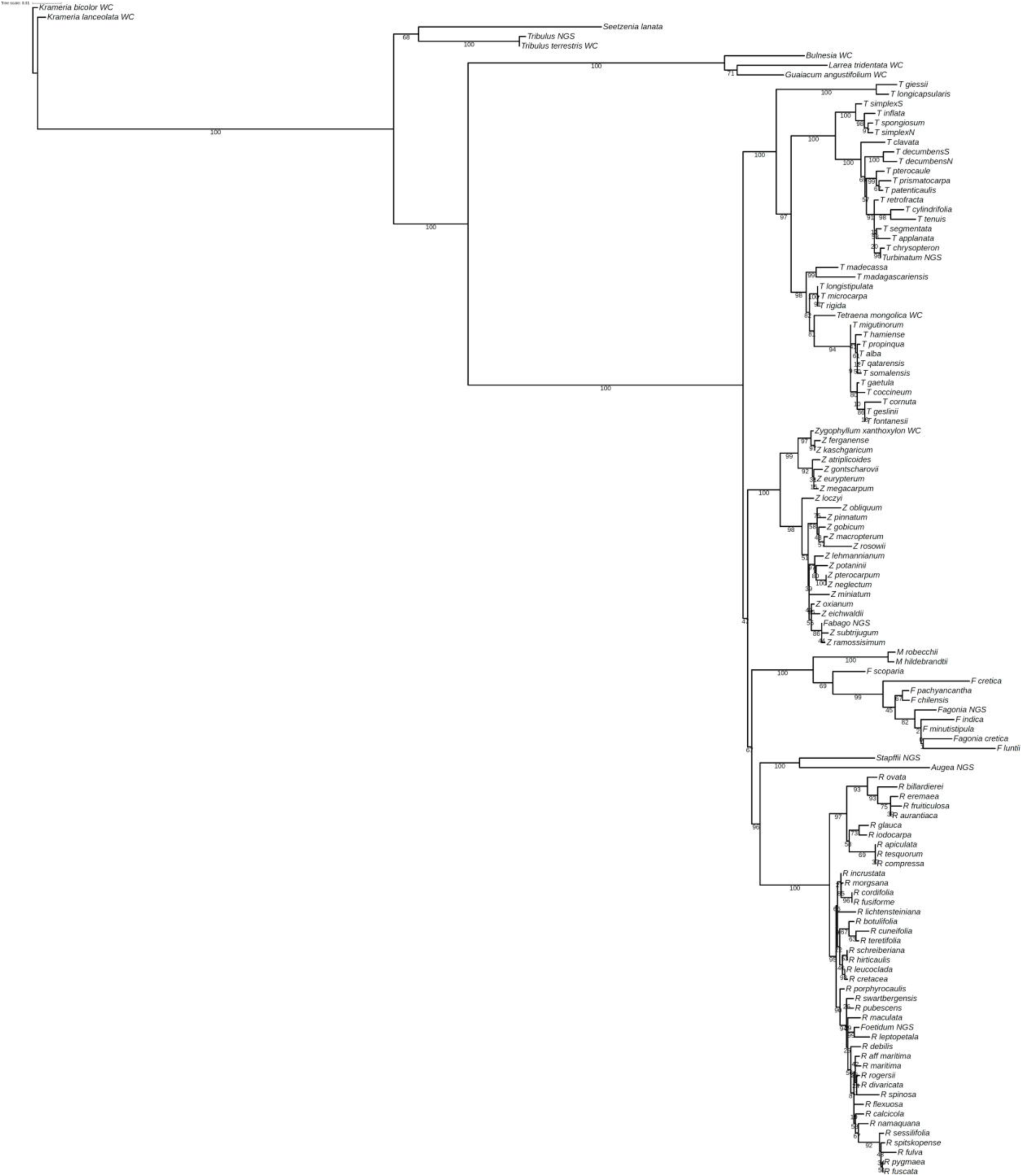
The most likely phylogenetic tree from the maximum likelihood analysis of the 3 HVNC genes plus 5 NC gene alignment matrix of 9 ingroup taxa and 8 outgroups plus 105 species = 122 taxa. Maximum likelihood bootstrap values are indicated on the nodes.

**Figure S6:**
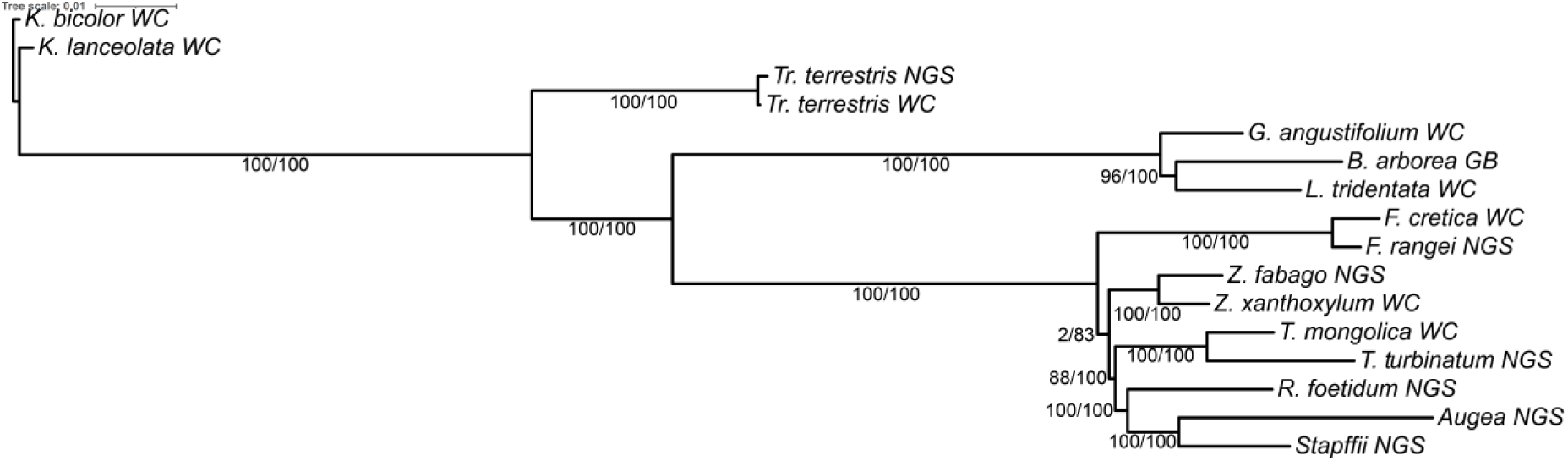
The most likely phylogenetic tree from the maximum likelihood analysis of the 16 taxa 18 coding genes plus 5 NC gene and 3 HVNC gene alignment matrix. Maximum likelihood bootstrap values followed by Bayesian posterior probabilities are indicated on the nodes.

**Figure S7:**
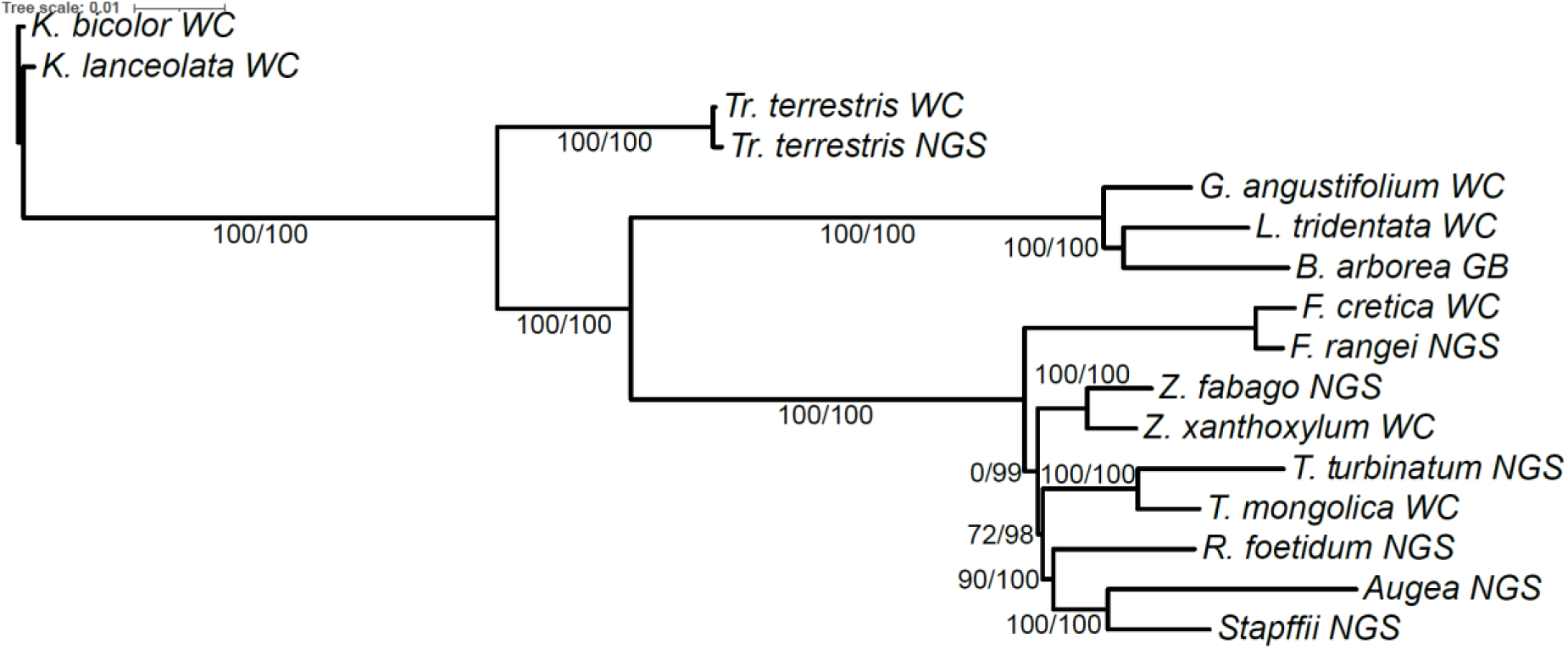
The most likely phylogenetic tree from the maximum likelihood analysis of the 16 taxa 39 coding genes plus 5 NC gene and 3 HVNC gene alignment matrix. Maximum likelihood bootstrap values followed by Bayesian posterior probabilities are indicated on the nodes.

**Figure S8:**
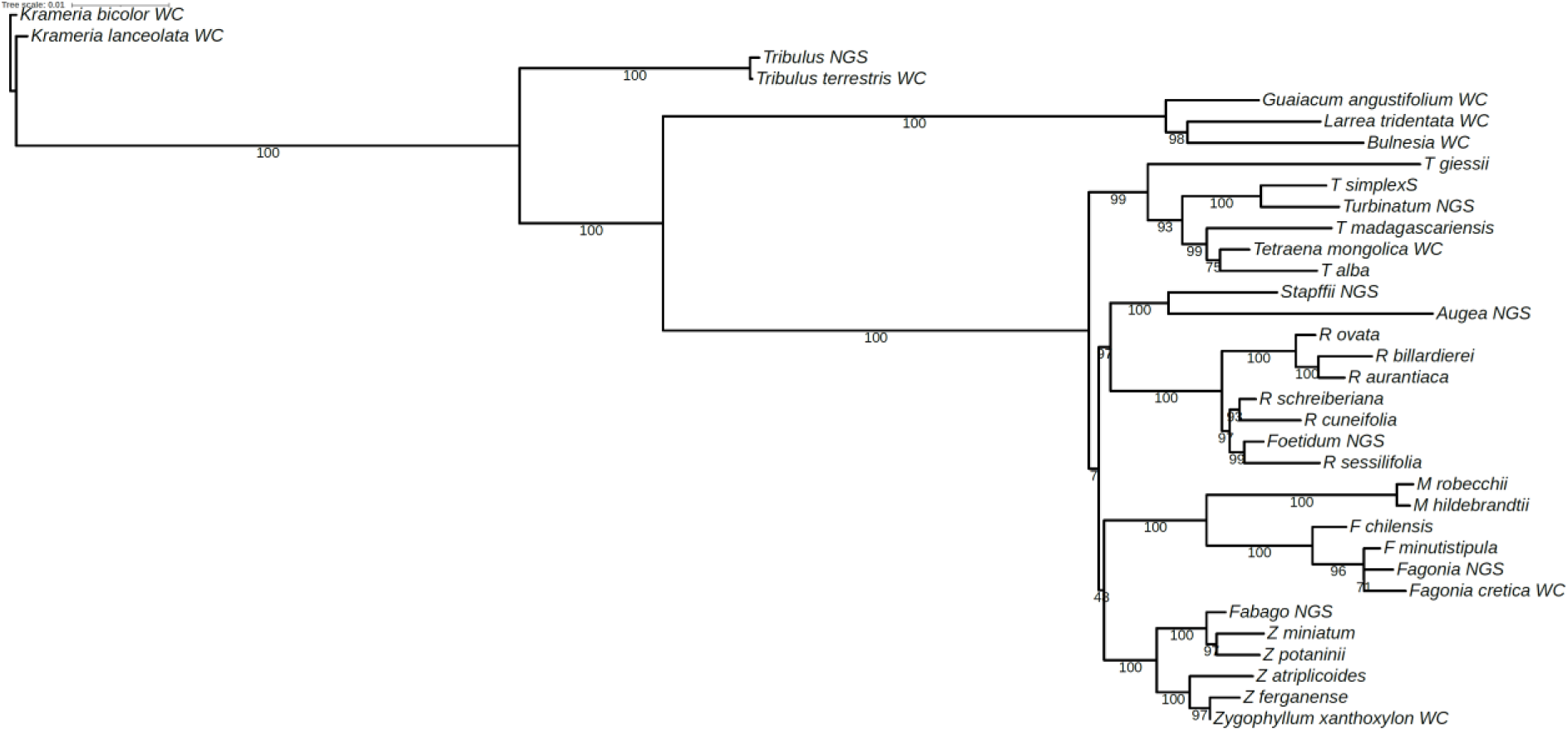
The most likely phylogenetic tree from the maximum likelihood analysis of the 3 HVNC genes plus 5 NC gene plus 39 coding gene alignment matrix of 9 ingroup taxa and 7 outgroups plus 18 species = 34 taxa. Maximum likelihood bootstrap values are indicated on the nodes.

**Figure S9:**
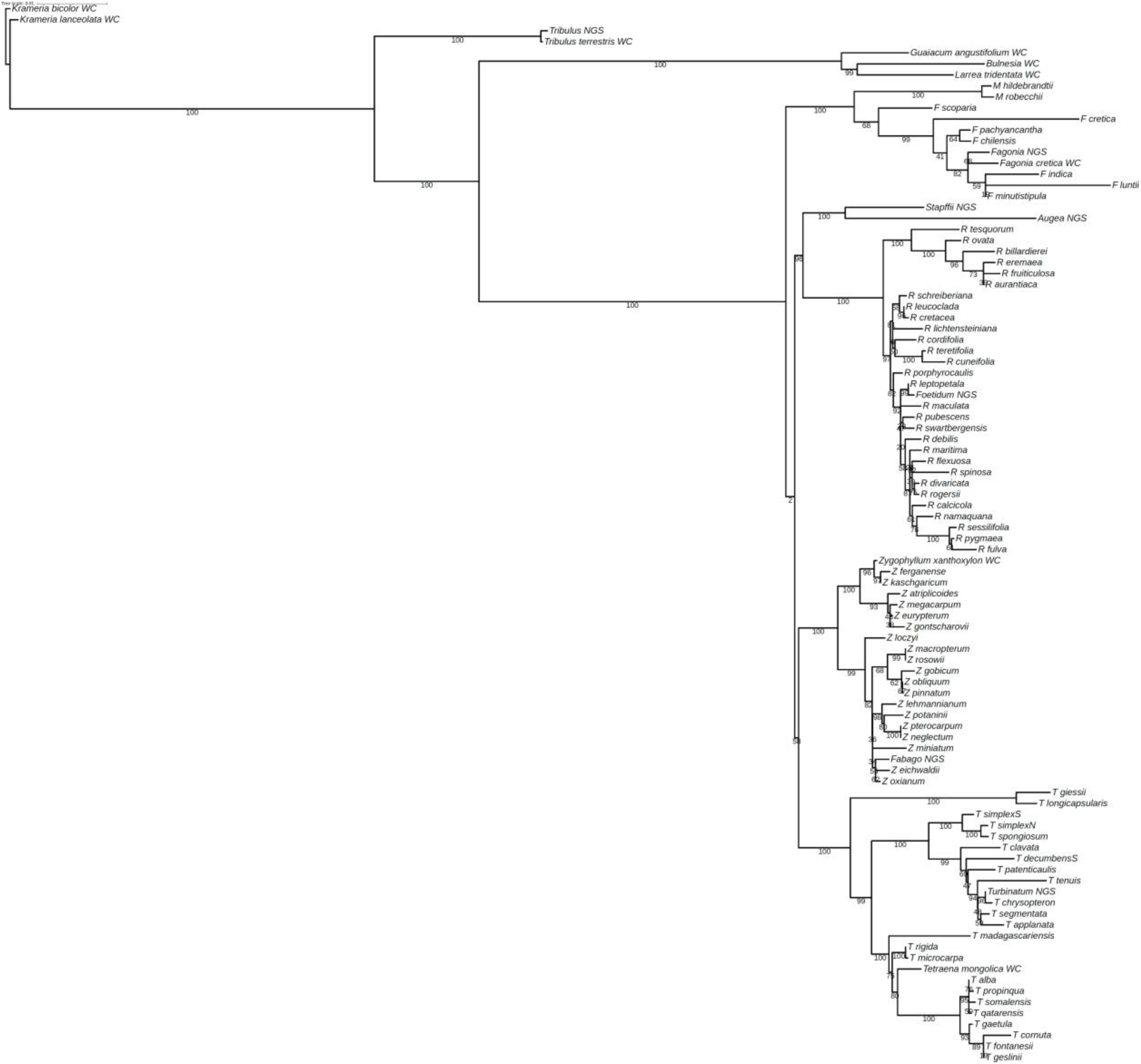
The most likely phylogenetic tree from the maximum likelihood analysis of the 3 HVNC genes plus 5 NC gene plus 39 coding gene alignment matrix of 9 ingroup taxa and 7 outgroups plus 80 species = 96 taxa. Maximum likelihood bootstrap values are indicated on the nodes.

**Figure S10:**
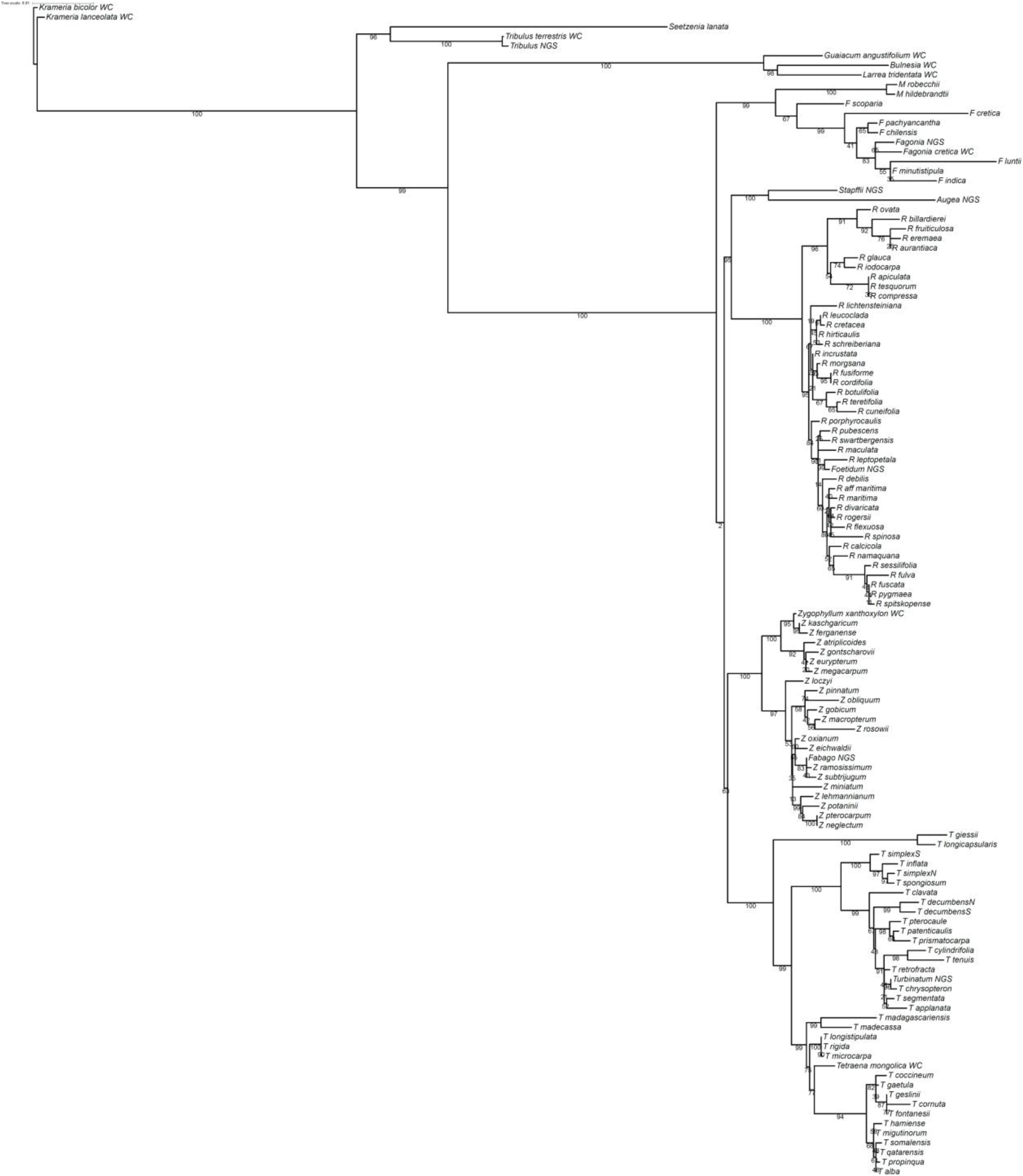
The most likely phylogenetic tree from the maximum likelihood analysis of the 3 HVNC genes plus 5 NC gene plus 39 coding gene alignment matrix of 9 ingroup taxa and 8 outgroups plus 105 species = 122 taxa. Maximum likelihood bootstrap values are indicated on the nodes.

**Figure S11:**
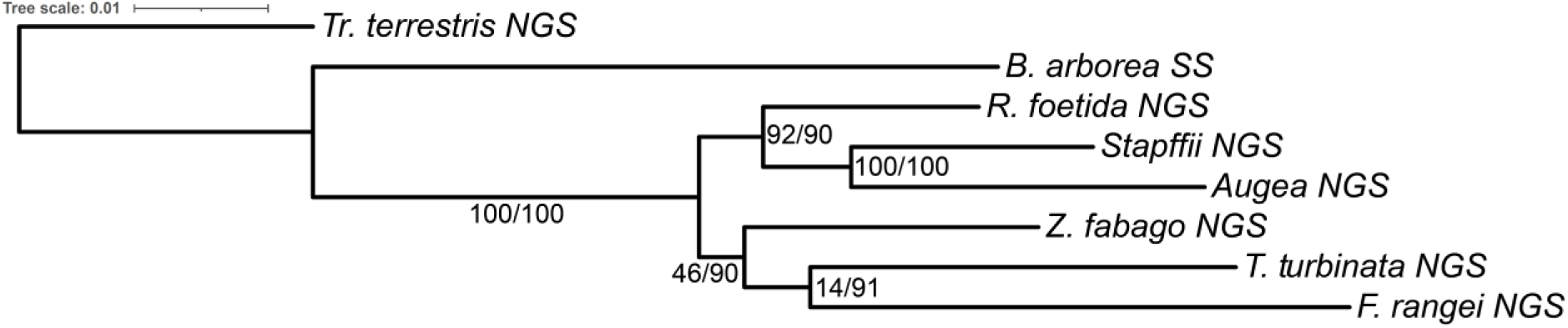
The most likely phylogenetic tree from the maximum likelihood analysis of the 8 taxa ITS cassette gene alignment matrix. Maximum likelihood bootstrap values followed by Bayesian posterior probabilities are indicated on the nodes.

**Figure S12:**
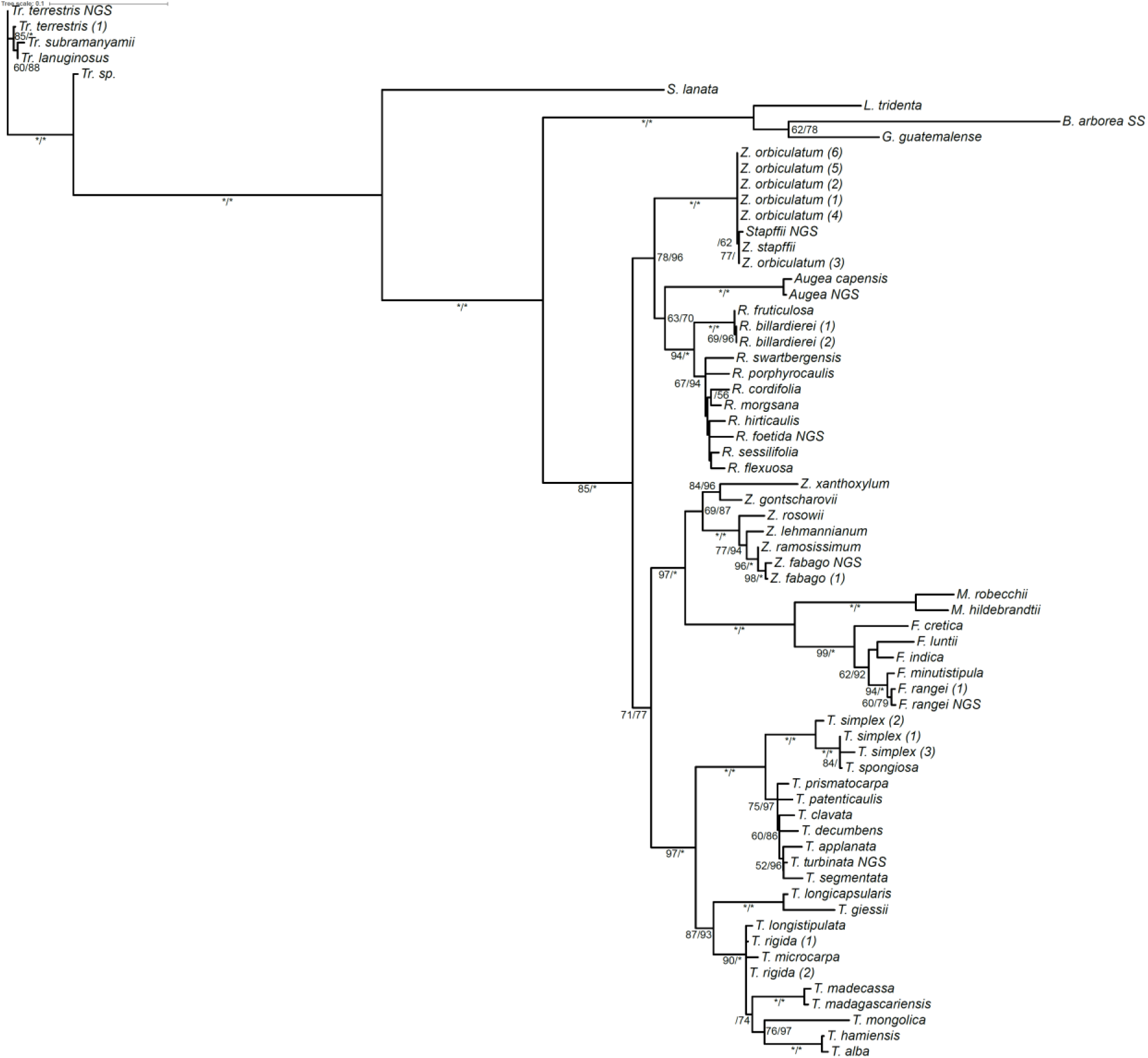
The most likely phylogenetic tree from the maximum likelihood analysis of the 67 taxa ITS region alignment matrix. Maximum likelihood bootstrap values followed by Bayesian posterior probabilities are indicated on the nodes. Where node support is 100 it is indicated as an asterisk.

